# Androgen and glucocorticoid receptor direct distinct transcriptional programs by receptor-specific and shared DNA binding sites

**DOI:** 10.1101/2020.10.15.340877

**Authors:** Marina Borschiwer, Melissa Bothe, Gözde Kibar, Alisa Fuchs, Stefanie Schöne, Stefan Prekovic, Isabel Mayayo Peralta, Ho-Ryun Chung, Wilbert Zwart, Christine Helsen, Frank Claessens, Sebastiaan H. Meijsing

**Author notes:** These authors contributed equally to this work (Marina Borschiwer, Melissa Bothe). Corresponding author, Contact information: Sebastiaan H. Meijsing, Max Planck Unit for the Science of Pathogens, Charitéplatz 1, Germany.

## Abstract

The glucocorticoid (GR) and androgen (AR) receptors execute unique functions *in vivo*, yet have nearly identical DNA binding specificities. To identify mechanisms that facilitate functional diversification among these transcription factor paralogs, we studied AR and GR in an equivalent cellular context. Analysis of chromatin and sequence features suggest that divergent binding, and corresponding gene regulation, are driven by different abilities of AR and GR to interact with relatively inaccessible chromatin. Divergent genomic binding patterns can also be the results of subtle differences in DNA binding preference between AR and GR. Furthermore, the sequence composition of large regions (>10 kb) surrounding selectively occupied binding sites differs significantly, indicating a role for the sequence environment in selectively guiding AR and GR to distinct binding sites. The comparison of binding sites that are shared between AR and GR shows that the specificity paradox can also be resolved by differences in the events that occur downstream of receptor binding. Specifically, we find that shared binding sites display receptor-specific enhancer activity, cofactor recruitment and changes in histone modifications. Genomic deletion of shared binding sites demonstrates their contribution to directing receptor-specific gene regulation. Together, these data suggest that differences in genomic occupancy as well as divergence in the events that occur downstream of receptor binding direct functional diversification among transcription factor paralogs.

## Introduction

The interplay between transcription factors (TFs), genomic DNA binding sites and the chromatin context in which recognition sequences are embedded plays a pivotal role in specifying where, when, and at which level genes are expressed. Differences in the DNA binding specificity among TFs can guide them to distinct genomic loci and allow TFs to carry out unique functions by regulating unique sets of target genes (Khan et al. 2018). However, even related TFs with very similar DNA-binding domains-and consequently overlapping DNA sequence preferences-perform non-redundant functions (Kribelbauer et al. 2019). Mechanistically, unique functions can derive from cell type-specific expression of related TFs (Singh and Hannenhalli 2008). However, functional diversification is also observed for paralogs that are expressed in the same cell type yet direct divergent genome-wide occupancy and gene regulation (Sahu et al. 2013; Jerković et al. 2017). One explanation for this is that subtle differences in the intrinsic DNA binding specificity among paralogs *in vitro* contributes to their differential binding *in vivo*. This was shown for paralogous TFs from different families with divergent binding mainly occurring at medium- and low-affinity binding sites (Shen et al. 2018; Zhang et al. 2018). The differences in sequence preference can occur within the core binding site but also in the regions flanking the motif (Levo et al. 2015; Gordân et al. 2013). Another explanation for functional diversification among paralogs was offered in a study of HOX proteins which showed that paralogs acquire novel and distinct DNA properties when they pair with another TF (Slattery et al. 2011). Of note, differential genome-wide occupancy can also be a consequence of different abilities to interact with relatively inaccessible chromatin as shown for Hox paralogs and for members of the Pou famility of TFs (Bulajić et al. 2019; De Kumar et al. 2017; Wapinski et al. 2013; Soufi et al. 2012). Finally, specificity could also be derived from events that occur downstream of binding when protein sequence differences between paralogs influence their activity. In this scenario, genomic binding sites that are shared, would selectively allow one of the TF paralogs to regulate the expression of target genes. Mechanistically, TF-specific activity from shared binding sites can be due to selective recruitment of cofactors, either coactivators or corepressors, that modulate transcriptional output (Joshi et al. 2010). However, the degree to which shared binding sites actually contribute to directing TF-specific gene regulation and the underlying mechanisms remain largely unexplored.

Steroid receptors are a family of ligand-dependent TFs and provide an attractive model to study functional diversification among paralogs given that their activity can be switched on or off by the addition or removal of their ligand. This on/off switch facilitates the relatively straightforward identification of changes in the cell, *e.g.* in gene expression, genome-wide binding and the chromatin landscape, that occur upon their activation. Two members of the steroid receptor family are the androgen receptor (AR) and the glucocorticoid receptor (GR). Despite their nearly identical DNA binding interface, the physiological roles of the hormones that activate AR (androgen) and GR (glucocorticoids), are different (Claessens et al. 2017). Androgens are male sex steroids and are, among other things, involved in development and the maintenance of reproductive organs. Glucocorticoids were named after their role in glucose metabolism as they promote gluconeogenesis in the liver and inhibit glucose uptake by skeletal muscle by antagonizing insulin but serve many additional functions. The different functions of AR and GR also translate into differences in their therapeutic use. Glucocorticoids are widely used to treat chronic inflammatory conditions with long-term use associated with muscle and bone wasting as side-effects (Klein 2015). In contrast, androgens have anabolic properties and can be used to treat glucocorticoid-induced osteoporosis showing that AR and GR can have antagonizing effects (Fraser and Adachi 2009). However, in certain castration-resistant prostate cancers, GR appears capable of taking over the role of AR as a driver of cancer progression, indicating the function of both receptors can also overlap depending on the context (Arora et al. 2013).

In line with both shared and divergent functions of AR and GR, their genome-wide binding patterns partially overlap in prostate cancel cells (Arora et al. 2013; Sahu et al. 2013). Accordingly, the genes regulated by AR and GR show some overlap but also diverge with each receptor also regulating a unique set of genes (Sahu et al. 2013; Arora et al. 2013). Shared genomic binding sites are expected given that AR and GR have nearly identical DNA binding interfaces and consensus sequence recognition motifs (Zhang et al. 2018). The mechanisms that direct differential binding and gene regulation by AR and GR are less clear. *In vitro* studies indicate that the intrinsic DNA binding preference varies between AR and GR with differences both within the core motif and in the sequences directly flanking it (Zhang et al. 2018). In addition, AR is able to selectively bind response elements that diverge from the consensus motif which consists of inverted hexameric half-sites separated by a 3 bp spacer. Selective AR binding sequences have a more degenerate second half-site, which often resembles a direct rather than an inverted repeat (Claessens et al. 2017). Together, these studies demonstrate that divergent DNA-binding preferences contribute to the differential binding observed *in vivo*, however this does not explain all differential binding observed *in vivo* (Sahu et al. 2013; Zhang et al. 2018). Moreover, selective binding does not explain how genes with nearby binding sites that are occupied by both AR and GR can be regulated in a receptor-specific manner (Arora et al. 2013).

Here, we set out to study the mechanisms that endow AR and GR with unique functions by examining human osteosarcoma cell lines that express either GR or AR. Specifically, we studied the role of chromatin and sequence in directing AR and GR to distinct genomic loci. Our results indicate that differences in binding can be explained by distinct DNA sequence preferences both within the motif and in the sequence composition of regions of several kb that surround sites that are selectively occupied by one of the receptors. In addition, we find indications that divergent binding is generated by receptor-specific abilities to interact with “inaccessible” chromatin. Finally, we compared receptor-induced changes that occur downstream of binding and find that binding sites that are shared by both receptors can nevertheless direct receptor-specific changes in histone modifications, enhancer activity and gene regulation. Together, our results indicate that divergence in gene regulation by AR and GR is driven by both difference in genomic binding and in the events that occur downstream of binding.

## Materials and Methods

### Experimental

#### Plasmids

The AR expression construct was modified from the pGFP-AR plasmid (Farla et al. 2004) by replacing a NheI-BspTI fragment containing *EGFP* and 510 bp of human *AR* and replacing it with a PCR-amplified fragment with primer-introduced ATG, NheI and BspTI sites. Individual STARR reporter constructs were generated by digesting the human STARR-seq vector (Arnold et al. 2013) with SalI and AgeI and subsequent insertion of fragments of interest by In-Fusion HD cloning (TaKaRa). Fragments of interest: positive control region (near *IP6K3* gene, hg19: chr6:33,698,504-33,698,853), *AQP3* enhancer (hg19: chr9:33,437,258-33,437,811) and the *AQP3* enhancer with AR binding site mutations (*AQP3*-Deleted and *AQP3*-AGA -> TGT) were ordered as a gBlock (IDT) or GeneStrand (Eurofins) (see Table 1 for the sequence of regions).

**Table 1:**
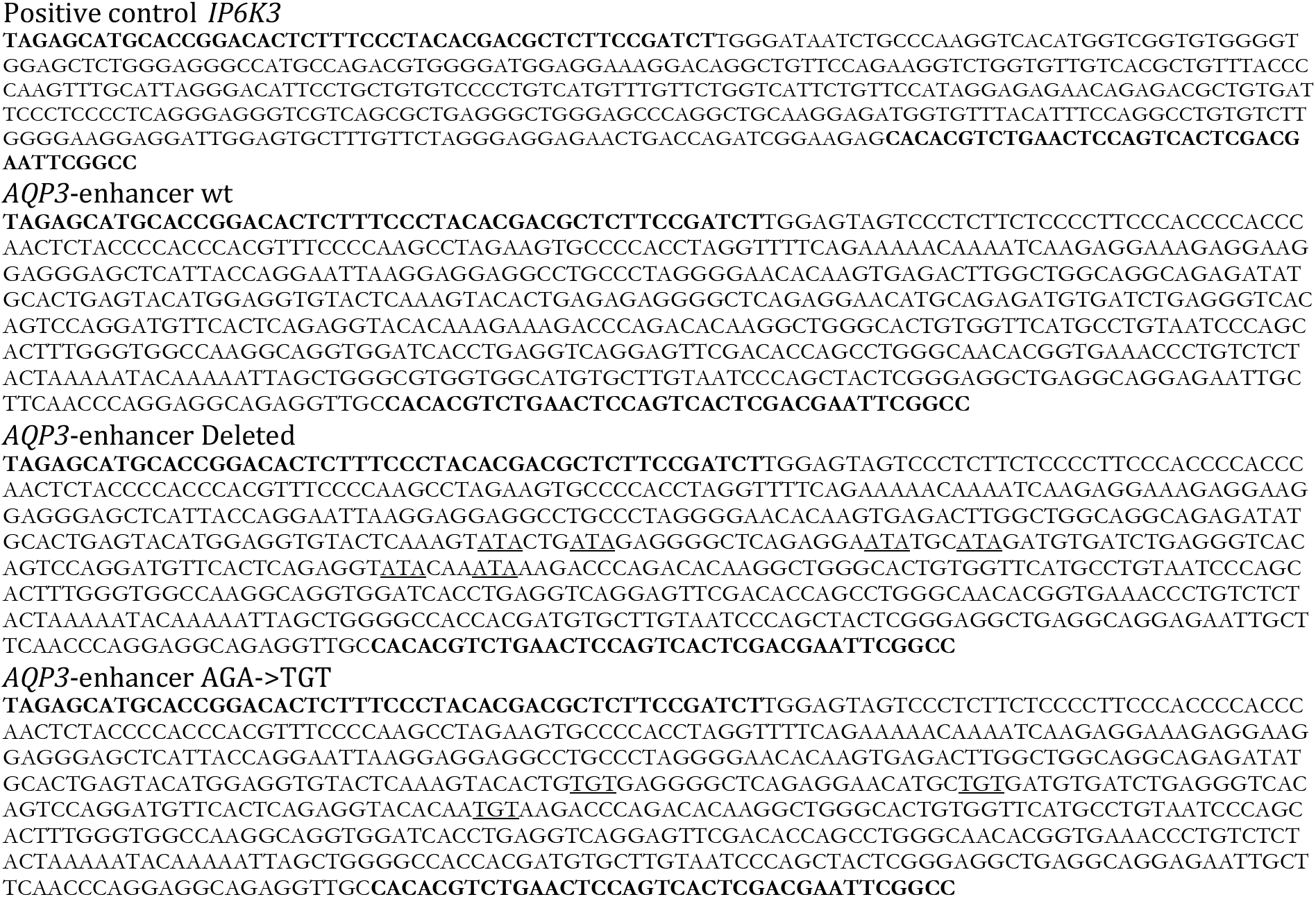
Regulatory regions of individual STARR-seq constructs.

Oligos (Table 2) encoding guide RNAs to delete the region downstream of the *AQP3* gene were designed using the CRISPR Design tool (Hsu et al. 2013), annealed and cloned into BbsI digested PX459 plasmid (Addgene #62988, (Ran et al. 2013)) to generate plasmids PX459-AQP3_214 and PX459-AQP3_216.

**Table 2:**
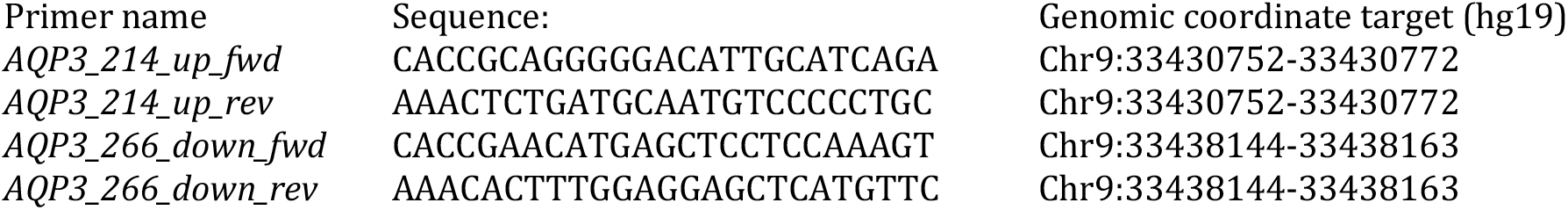
Primers to clone sgRNAs targeting the AQP3 locus.

#### Cell lines, Transient transfections

U2OS cells stably transfected with rat GRα (U2OS-GR) (Rogatsky et al. 1997) were grown in DMEM supplemented with 5% FBS. U2OS cells stably transfected with human AR (U2OS-AR) were generated as follows: U2OS cells were transfected with 30 ng of the AR expression construct using Lipofectamine and Plus Reagent (Invitrogen). The next day, ~10.000 cells were transferred to a 15 cm dish and single-cell derived clonal lines stably transfected with the AR expression construct were selected using G418 (800 μg/ml).

To test the activity of individual STARR reporters, 1 million U2OS cells were transfected using the Nucleofector 2b with 2 μg Plasmid DNA using kit V (Lonza) according the manufacturer’s instructions.

#### Whole-cell [^3^H] steroid binding assay

Hormone binding assays were performed essentially as described (Meijsing et al. 2007). In brief, 25.000 cells (U2OS-GR or U2OS-AR) were seeded per well of a 24-well plate. One day before the assay, the medium was replaced with DMEM containing 5 % charcoal-stripped FBS. U2OS-GR cells were treated with 100 nM [^3^H-]Dexamethasone (Dex) (81 Ci/mmol, PerkinElmer) in the presence or absence of a 10 μM (100 x) excess of unlabeled Dex for 45 min. U2OS-AR were treated with 100 nM [^3^H]-R1881 (81.4 Ci/mmol, PerkinElmer) in the presence or absence of a 10 μM (100 x) excess of unlabeled R1881 for 45 min. After five washes with ice-cold PBS, the ligand was extracted by adding 250 μl of ethanol for 45 min and quantified by liquid scintillation counting. Specific activity of [^3^H]-Dex and [^3^H]-R1881 and the number of cells per well were used to calculate the number of bound [^3^H] molecules per cell. Specific steroid binding was calculated as the difference between total and nonspecific binding.

#### Immunoblotting

Total protein from equal amounts of cells was separated on a NuPage Gradient 4-12% Bis-Tris Mini Gel (Invitrogen), transferred to a PVDF membrane, and incubated with either anti-AR (Sigma Aldrich, 06-680, 1:1000) or anti-actin (Sc-1616; Santa Cruz Biotechnology, 1:1000) antibodies followed by incubation with secondary antibodies conjugated with horseradish peroxidase (Invitrogen, 656120, 1:4000). Proteins were visualized using the SuperSignal West Dura Extended Duration Substrate (ThermoFisher).

#### RNA-seq

For U2OS-GR, the RNA-seq data was from previous studies (Thormann et al. 2019; Schöne et al. 2018), ArrayExpress accession number E-MTAB-6738. U2OS-GR cells were treated for 4h with either 1μM Dex or 0.1% ethanol as vehicle control. For U2OS-AR, cells were treated for 24h with either 5 nM R1881 or 0.1 % dmso as vehicle control. mRNA was isolated from 1.2 million cells using the RNeasy kit from Qiagen and oligo d(T) beads. Sequencing Libraries were prepared for paired end Illumina sequencing using the TruSeq RNA library Prep Kit (Illumina). ArrayExpress accession number: E-MTAB-9622.

#### RNA Isolation, cDNA Synthesis and qPCR analysis

RNA was isolated using a RNeasy Mini Kit (Qiagen). For the analysis of cells transfected with STARR reporters, total RNA was reverse-transcribed with the PrimeScript One Step Kit (Takara) using gene-specific primers for *GFP* (CAAACTCATCAATGTATCTTATCATG) and *RPL19* (GAGGCCAGTATGTACAGACAAAGTGG) which was used for data normalization. qPCR and data analysis were done as described (Meijsing et al. 2009). For all other experiments, cDNA synthesis was performed with Random Primer and Oligo dT primer provided by the PrimeScript One Step Kit. Primers for qPCR are listed in Table 3.

**Table 3:**
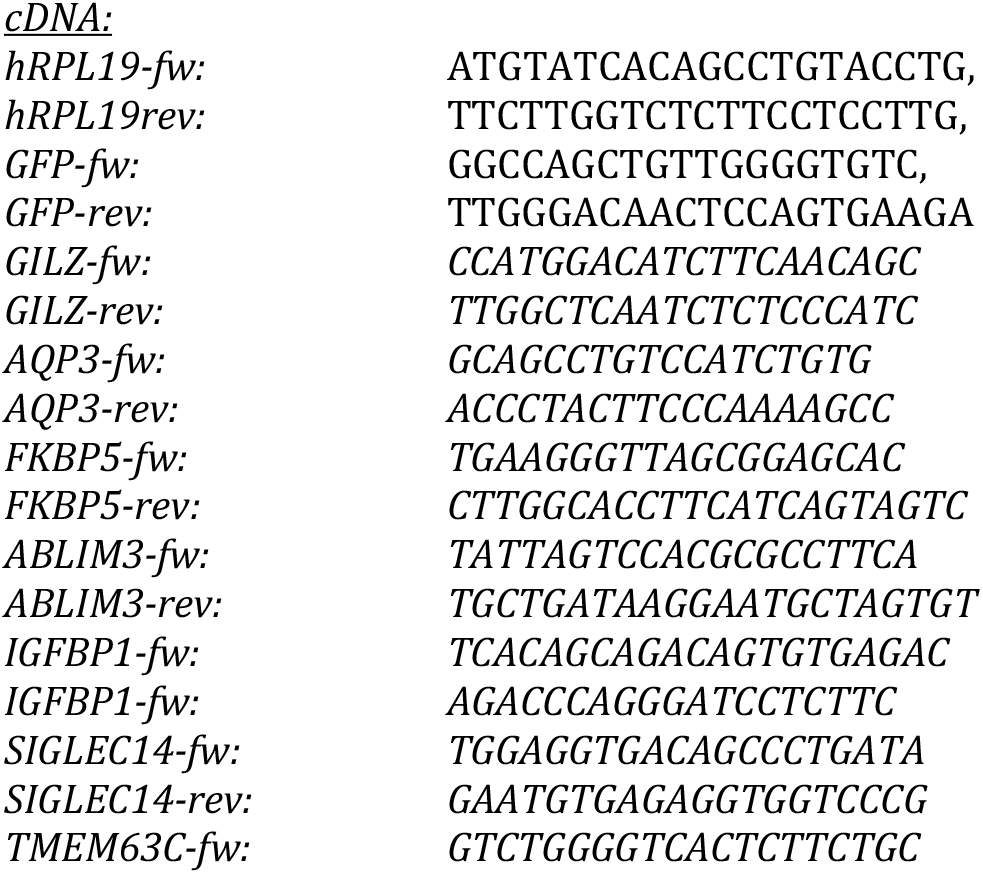
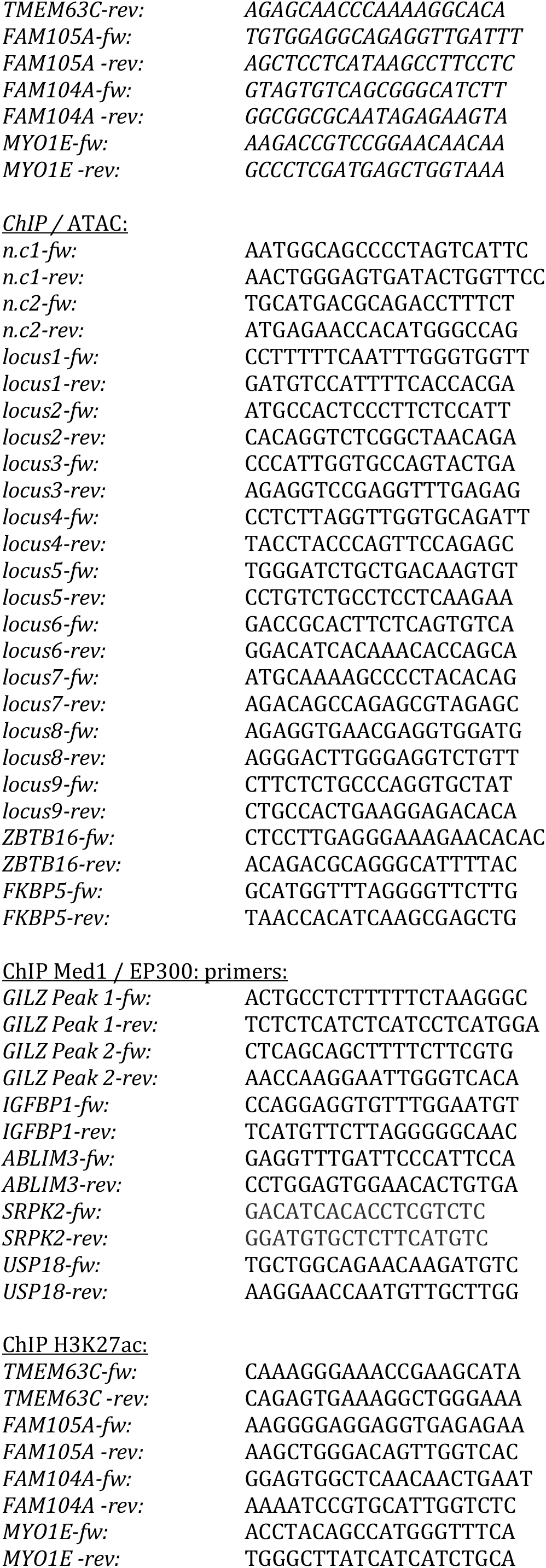
Primer pairs for qPCR:

#### ChIP-seq and ChIP-qPCR

ChIP assays were performed using the following antibodies: GR, N499 (2 μl/ChIP); AR, polyclonal antibody PG-21 (Anti-AR; Sigma Aldrich; 06-680, 2 μl/ChIP); H3K27ac, C15410196 (Diagenode 1μg /ChIP); Med1, A300-793A (Bethyl Laboratories, 5 μl/ChIP); EP300, C15200211 (Diagenode, 5 μl/ChIP); H3K4me1, C15410194 (Diagenode 1μg/ChIP); H3K9me3, C15410193 (Diagenode 1μg/ChIP); H3K27me3, C1541095 (Diagenode 1μg/ChIP).

ChIP-seq data for GR replicate 1 (1 μM Dex, 1.5h) is from SRA accession number SRP020242 (Schiller et al. 2014). ChIP-seq for GR replicate 2 (1 μM Dex, 1.5h) and ChIP-seq for AR (5 nM R1881, 4h) was done as described (Meijsing et al. 2009). ChIP-seq experiments targeting histone marks in U2OS-AR (5 nM R1881 or 0,1 % dmso as a vehicle control, (4h) and U2OS-GR18 (1 μM Dex or 0.1 % EtOH as vehicle control, 1.5h) cells were done as described before for H3K27ac (Thormann et al. 2018). Sequencing libraries were prepared using the NEBNext Ultra DNA Library Prep kit (E7370; NEB) according to the manufacturer’s instructions and submitted for paired-end Illumina sequencing. ArrayExpress accession numbers for ChIP-seq data we generated: E-MTAB-9616 and E-MTAB-9617.

ChIP assays for subsequent analysis by qPCR were done as described (Meijsing et al. 2009) with the treatment times and hormone concentrations as described above. For Med1 and EP300 ChIPs, cells were cross-linked with 1% formaldehyde for 10 min instead of 3 min as used for all other targets. Hormone treatment was 1.5h for both AR and GR. Primers for qPCR are listed in Table 3.

#### ATAC-seq and ATAC-qPCR

ATAC-seq data for GR (1 μM Dex, 1.5h or 0.1 % EtOH as vehicle control) is from a previous study (Thormann et al. 2019), ArrayExpress accession number E-MTAB-7746.

For AR (5 nM R1881 or 0.1 % dmso as a vehicle control, 4h), ATAC-seq was done as described (Thormann et al. 2019). ArrayExpress accession number for ATAC-seq data: E-MTAB-9606.

For the analysis of ATAC experiments by qPCR, DNA fragments were amplified using p5 and p7 primers, two-sided size selected using AMPure XP beads (Beckman Coulter) and analyzed by qPCR using the primers listed in Table 3.

#### FAIRE-STARR-seq

##### Library construction

Accessible chromatin regions were isolated from U2OS-GR cells treated for 1.5h with 1 μM Dex, using the FAIRE method (Simon et al. 2013). In short, cells were fixed with 1% formaldehyde, DNA was isolated from the aqueous phase, cross-linking was reversed, and the DNA was purified. The resulting library of DNA fragments was ligated to Illumina adapters (NEB #E7335) using the NEBNext Ultra DNA Library kit (NEB #E7370) according to manufacturer’s instructions, except that the final PCR amplification step was omitted. Instead, six PCR reactions with 2μl adapter-ligated DNA were preformed using NEBNext Q5 Hot Start HiFi PCR Master Mix and primers with 15nt extension for subsequent In-Fusion cloning (fw: TAGAGCATGCACCGGACACTCTTTCCCTACACGACGCTCTTCCGATCT & rev: GGCCGAATTCGTCGAGTGACTGGAGTTCAGACGTGTGCTCTTCCGATCT) (Arnold et al. 2013). The resulting DNA fragments were cloned into the linearized STARR-seq vector (Addgene #71509, *AgeI*-*SalI* digested) using the In-Fusion HD kit (Takara), followed by transformations into MegaX DH10B T1R electrocompetent cells (ThermoFisher Scientific). A total of 20 In-Fusion HD reactions and transformations were performed, pooled and the plasmid library was extracted using a Plasmid Plus Mega Kit (Qiagen). *Transfection of U2OS-AR or U2OS-GR cells.* 5 million cells were transfected with 5 μg of DNA FAIRE-STARR library plasmid using the Amaxa Nucleofector kit V (Lonza). For each condition, 4 transfections were pooled and seeded into two 15cm dishes with medium containing PKR (C16, Sigma; cat# I9785-5MG) and TBK1/IKK inhibitors (BX-795, Sigma; cat# SML0694-5MG) at a final concentration of 0.5 μM per inhibitor to suppress the type I interferon response (Muerdter et al. 2018). U2OS-GR cells were treated with 1 μM Dex or 0.1 % EtOH as a vehicle control. U2OS-AR cells were treated with 5 nM R1881 or 0.1 % dmso as vehicle control. After 14h, cells were harvested by trypsinization, snap frozen and stored at −80°C.

##### FAIRE-STARR-seq library preparation

RNA isolation, reverse transcription and amplification of cDNA for subsequent Illumina 50bp paired-end sequencing were essentially done as described (Arnold et al. 2013) except that a modified primer (5’-CAAGCAGAAGACGGCATACGAGATnnnnnnnnGTGACTGGAGTTCAGACGTGTGCTCTTCCGA

TCT −3’) was used during the reverse transcription step to introduce unique molecular identifiers (UMIs). 50 ng cDNA/reaction was used as a template in every 50 μL PCR reaction using Kapa Hifi hotstart ready mix (Roche). PCR conditions: 98 °C for 45 s; 15 cycles of 98 °C for 15 s, 65 °C for 30 s and 72 °C for 30 s. Illumina HiSeq-compatible primers were used (forward: 5’-CAAGCAGAAGACGGCATACGA-3’; reverse: 5’-AATGATACGGCGACCACCGAGATCTACAC-index-ACACTCTTTCCCTACACGACGCTC-3’).

PCR products were purified with AMPure XP beads (Beckman Coulter; beads/reaction ratio = 1) and submitted for paired-end Illumina sequencing. ArrayExpress accession number for STARR-seq data: E-MTAB-9614.

#### Genome editing using CRISPR/Cas9

Clonal lines with CRISPR/Cas9-deleted GR-bound regulatory regions near the *GILZ* gene (*GILZ ΔGBS1-4, ΔGBS A-G*) were from a previous study (Thormann et al. 2018). To generate single-cell-derived clonal lines lacking an AR-bound region upstream of the *AQP3* gene, 1 million U2OS-AR cells were transfected with 1.2 μg each of the PX459-AQP3_214 and PX459-AQP3_216 plasmids using the Amaxa V Kit (Lonza). Cells were plated and the next day we selected for transfected cells by refeeding cells with medium containing puromycin (10 μg/ml). 24h later, surviving cells were trypsinized and seeded to isolate single-cell-derived clonal lines. To genotype single cell-derived clonal lines, genomic DNA was isolated using the Blood and Tissue kit (Qiagen), the targeted region was amplified by PCR using primers spanning the target region (Table 4) and deletion of the region was analyzed by agarose gel electrophoresis and Sanger sequencing. The residual presence of wt alleles was analyzed using primers spanning the breakpoints at both ends of the targeted locus (Table 4).

#### RIME

After hormone deprivation, the U2OS-GR and U2OS-AR cell lines were treated with either 1 μM Dex or 5 nM R1881 for 4h, respectively. Next, RIME experiments were performed and analyzed as previously described (Mohammed et al. 2016). The following antibodies were used: anti-GR (12041, Cell Signalling Technology), anti-AR (06-680, Merck), or anti-rabbit IgG (sc-2027, Santa Cruz Biotechnology) as control. Tryptic digestion of bead-bound proteins was performed as described previously (Stelloo et al. 2018). LC-MS/MS analysis of the tryptic digests was performed on an Orbitrap Fusion Tribrid mass spectrometer equipped with a Proxeon nLC1000 system (Thermo Scientific) using the same settings, with the exception that the samples were eluted from the analytical column in a 90-min linear gradient. Raw data were analyzed by Proteome Discoverer (PD) (version 2.3.0.523, Thermo Scientific) using standard settings. MS/MS data were searched against the Swissprot database (released 2018_06) using Mascot (version 2.6.1, Matrix Science, UK) with *Homo sapiens* as taxonomy filter (20,381 entries). The maximum allowed precursor mass tolerance was 50 ppm and 0,6 Da for fragment ion masses. Trypsin was chosen as cleavage specificity allowing two missed cleavages. Carbamidomethylation (C) was set as a fixed modification, while oxidation (M) and deamidation (NQ) were used as variable modifications. False discovery rates for peptide and protein identification were set to 1% and as additional filter Mascot peptide ion score>20 or Sequest HT XCorr>1 was set. The PD output file containing the abundances was loaded into Perseus (version 1.6.1.3) LFQ intensities were Log2-transformed and the proteins were filtered for at least 66% valid values. Missing values were replaced by imputation based on the standard settings of Perseus, i.e. a normal distribution using a width of 0,3 and a downshift of 1,8. Differentially enriched proteins were called using a t-test. Data corresponding to RIME experiments was plotted using ggplot2 package in R. The geneset enrichment analysis was performed using GSEA-R (https://github.com/GSEA-MSigDB/GSEA_R) (Subramanian et al. 2005; Mootha et al. 2003).

**Table 4:**
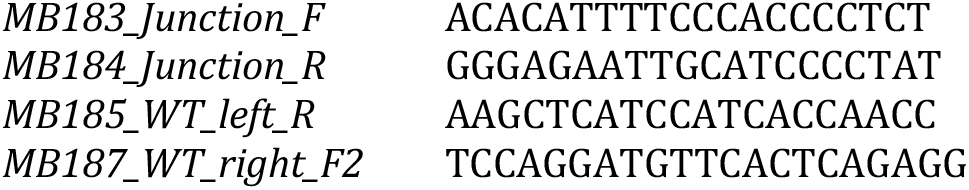
Primers to genotype AQP3 clonal lines.

### Computational

#### RNA-seq

Paired end 50bp reads from Illumina sequencing were mapped against the human hg19 reference genome using STAR (Dobin et al. 2013) (options: --alignIntronMin 20 -- alignIntronMax 500000 --chimSegmentMin 10 --outFilterMismatchNoverLmax 0.05 -- outFilterMatchNmin 10 --outFilterScoreMinOverLread 0 -- outFilterMatchNminOverLread 0 --outFilterMismatchNmax 10 -- outFilterMultimapNmax 5). Differential gene expression between hormone-treated and vehicle conditions from three biological replicates was calculated with DESeq2 (Love et al. 2014), default parameters except betaPrior=FALSE.

#### Overlap gene regulation by AR and GR (Venn diagram)

Differentially expressed genes (DEGs) between the hormone-treated samples and vehicle condition samples were identified both for AR (24h post hormone treatment vs control) and GR (4h post hormone treatment vs control) using the *LfcShrink* function in *DESeq2* R package (version 1.24.0, (Love et al. 2014)). Genes with an adjusted p-value < 0.05 and log2(fold change) > 1.5 were designated as significant upregulated genes. As a result, 777 and 364 upregulated genes (187 shared) were identified for GR and AR respectively.

#### ChIP-seq, ChIP BigWig tracks for genome browser screenshot

ChIP-seq reads were mapped with Bowtie2 v2.1.0 (Langmead and Salzberg 2012) (options: --very-sensitive) to the hg19 reference genome. The ChIP-seq reads for GR replicate 1 (SRP020242, (Schiller et al. 2014)) were mapped with Bowtie2 to hg19 (options: --very-sensitive -X 600 --trim5 5). Reads of mapping quality <10 were filtered out using SAMtools v1.10 (Li et al. 2009). Picard tools v.2.17.0 (http://broadinstitute.github.io/picard/) (MarkDuplicates) was used to remove duplicate reads. BigWig tracks for genome browser visualization were generated with deepTools v3.4.1 (Ramírez et al. 2014) bamCoverage (options: --normalizeUsing RPKM --binSize 20 --smoothLength 60). AR and GR ChIP-seq peaks for each replicate were called over their respective inputs using MACS2 v2.1.2 (Zhang et al. 2008) with a qvalue cut-off of 0.05. The final set of peaks was obtained by extracting overlapping peaks between both replicates using BEDtools intersect v2.27.1 (-u) (Quinlan and Hall 2010) and removing ENCODE blacklisted regions for hg19 (ENCODE Project Consortium 2012) as well as regions within unassigned contigs (chrUn) and mitochondrial genes (chrM). Shared and receptor-specific peaks were obtained with BEDtools intersect. GR-specific peaks at regions of low accessibility were obtained by sorting the ATAC-seq signal (vehicle control treatment) of U2OS-GR cells in descending order at all GR-specific peaks (+/− 250 bp around the peak center) using computeMatrix from deepTools and subsequently extracting the bottom 17,125. Peak calling was performed on ATAC-seq data (vehicle control treatment) in U2OS-AR cells and on ATAC-seq data (vehicle control treatment) in U2OS-GR cells using MACS2 v2.1.2 (Zhang et al. 2008) (options: --broad – broad-cutoff 0.05) and resulting peaks were removed from the GR-specific peaks with low accessibility, to ensure the list represented inaccessible sites.

#### ATAC-seq data processing

Processing of ATAC-seq data was done as previously described in (Thormann et al. 2019), with the addition that reads of mapping quality <10 were filtered out using SAMtools v1.10 (Li et al. 2009). BigWig tracks for genome browser visualization were generated with bamCoverage (options: --normalizeUsing RPKM; --binSize 20; -- smoothLength 60) from deepTools v3.4.1 (Ramírez et al. 2014).

#### FAIRE-STARR-seq data processing

FAIRE-STARR-seq reads were mapped with Bowtie2 v2.1.0 (Langmead and Salzberg 2012) (options: --very-sensitive) to the hg19 reference genome. Reads were deduplicated based on their UMIs and genomic coordinates using the UMI-tools v1.0.0 dedup function (Smith et al. 2017). SAMtools v1.10 (Li et al. 2009) was used to filter out reads of mapping quality <10. Replicate BAM files were merged with SAMtools merge for downstream analyses. For genome browser visualization, bigWig tracks of merged BAM files were generated with bamCoverage (options: --normalizeUsing RPKM; -- binSize 20; --smoothLength 60) from deepTools v3.4.1 (Ramírez et al. 2014).

#### Heatmaps and mean signal plots at AR and GR peaks

Heatmaps and their respective mean signal plots (+/− 2 kb around the peak center) of shared and receptor-specific peaks were generated with computeMatrix (options: reference-point) and plotHeatmap from deepTools v3.4.1 (Ramírez et al. 2014), using bigWig files as input which had been generated with deepTools bamCoverage (options: --normalizeUsing RPKM).

#### Intersect binding and gene regulation

##### Gene categories

Receptor-specific and non-regulated genes were defined as follows: receptor-specific: adjusted p-value < 0.05 and log2(fold change) > 1.5 for either AR or GR; Shared: adjusted p-value < 0.05 and log2(fold change) > 1.5 for both AR and GR; non-regulated: adjusted p-value < 0.5 and 0.5 > log2(fold change) > 0.

##### Assigning ChIP-seq peaks to genes

For each gene, we scanned a 60-kb genomic window centered on transcriptional start sites (TSSs) for overlap with each peak file (AR-specific peaks, GR-specific peaks, shared peaks between AR and GR) using bedtools window. TSS annotations of genes (hg19) were obtained from the NCBI RefSeq database using the SeqMiner package (Zhan and Liu 2015). To make sure that each peak is only assigned to one of the gene categories, peaks assigned to more than one gene category were assigned to the nearest gene using the package ChIPpeakAnno in R (Zhu et al. 2010). Genes lacking binding sites in the 60kb window are labelled as ‘no peaks’. The distribution of binding events for each gene category are shown as stacked bar graphs. We used the same approach to link GR-specific peaks in regions of low chromatin accessibility to the different gene categories.

Statistical tests were performed using the Fisher Exact test on 2×2 contingency tables comparing the number of genes in each differential gene category that have specific binding events to the number of non-regulated genes that have corresponding binding events.

#### FAIRE-STARR and other features at shared peaks near different gene categories

FAIRE-STARR-seq (window +/− 250 bp), H3K27ac, AR/GR ChIP-seq or ATAC-seq mean signal (window +/− 2 kb around the peak center) was plotted for shared peaks of the different gene categories (for categorization see: Intersect binding and gene regulation) using deepTools v3.4.1 (Ramírez et al. 2014) computeMatrix (options: reference-point) and plotProfile, using BigWigs files as input.

#### Motif analysis and GC content of AR and GR-specific binding sites, AR binding sites of the *AQP3* enhancer

Motif enrichment analysis at AR and GR binding sites was performed with AME (McLeay and Bailey 2010) of the MEME suite v5.1.1 (Bailey et al. 2009). All AR-specific peaks as well as GR-specific peaks (either 6593 peaks randomly sampled or the 6593 peaks with the highest chromatin accessibility as sorted by ATAC-seq signal (vehicle treatment) of U2OS-GR) were used as input. The peak sequences (+/− 250pb around the peak center) were scanned for the JASPAR 2018 CORE Vertebrates Clustering motifs (Khan et al. 2018) including the AR-specific DR3 motif (Fig. S4A). Control sequences were either set to be the shuffled input sequences or peak sequences (+/− 250 bp around the center) of the other hormone receptor. For the heatmap representation, motif hits were included if the E value was <10^−30^ for either AR or GR. Alternatively, the top 7 motif logos are shown.

GC content profiles were generated at all AR-specific and all GR-specific peaks as well as at high accessibility AR- and GR-specific peaks. The high accessibility regions were obtained by sorting the ATAC-seq signal (vehicle treatment) at all AR- or GR-specific peaks (+/− 250 bp around the peak center) of the respective cell line in descending order and extracting the top 3296 peaks. Next, the hg19 reference genome was binned into 50 bp bins with the BEDtools v2.27.1 (Quinlan and Hall 2010) makewindows function. For each bin, the GC content was obtained using BEDtools nuc and the resulting bedgraph file was converted into bigWig format with bedGraphToBigWig (Kent et al. 2010). The GC content profiles +/− 5 kb around peak centers were generated using the deepTools v3.4.1 (Ramírez et al. 2014) commands computeMatrix (options: reference-point) and plotProfile.

For GC content plots in VCaP and LNCaP-1F5 cells, processed AR (100 nM dihydrotestosterone, 2h) and GR (100 nM Dex, 2h) ChIP-seq peaks from a previous study (Sahu et al. 2013) were downloaded from GEO (GSM980657, GSM980658, GSM980660, GSM980662, GSM980664). For AR peaks in VCaP cells, peaks from both replicates were intersected using BEDtools intersect v2.27.1 (-u) (Quinlan and Hall 2010) and overlapping peaks were extracted. ENCODE blacklisted regions for hg19 (ENCODE Project Consortium 2012) were removed from all peaks. Shared and receptor-specific peaks for each cell line were obtained with BEDtools intersect. GC content profiles were plotted for all AR- and GR-specific regions in each cell line. Statistical tests were performed using Mann-Whitney-U test comparing GR and AR-specifically occupied regions. To identify motif matches in the *AQP3* enhancer sequence [hg19: chr9:33437258-33437811], it was scanned for the AR (JASPAR ID MA0007.1-3) and GR (JASPAR ID MA0113.1-3) motif using the JASPAR CORE database (Sandelin et al. 2004). A total of six putative AR and four GR sites were found with a relative profile score threshold 80%. The top three AR MA0007.1-3 motif hits are shown (Fig. 7B).

#### Exoprofiler profiles

The ExoProfiler package was used to generate footprint profiles (Starick et al. 2015). For AR profiles in LNCaP cells, published ChIP-exo data (GSE43791) (Chen et al. 2015), mapped to hg19 with Bowtie2 v2.1.0 (Langmead and Salzberg 2012) (options: --very-sensitive) and filtered for mapping quality >10 using SAMtools v1.10 (Li et al. 2009), and AR peaks obtained from published ChIP-seq data (GSE43791) (Chen et al. 2015) were used as input. For GR profiles in U2OS-GR cells, published ChIP-exo data (EBI ArrayExpress E-MTAB-2955) (Starick et al. 2015) and GR peak regions were used as input. Peak sequences were scanned for the JASPAR motifs MA0113.2 and MA0007.2 (Khan et al. 2018) as well as the DR3 motif (Fig. S4A). The p-value threshold for motif matches was <10^−4^.

#### Heatmaps shared, AR-specific, non-regulated genes, GR-specific RIME interacting genes (mediator and chromatin categories)

Using the function *normTransform* in *DESeq2* (Love et al. 2014), the un-normalized gene counts were transformed into log transformed normalized gene counts for heatmap visualization of genes from previously obtained differential gene categories. To check the gene expression of AR-specific, GR-specific upregulated genes after hormone treatment in AR and GR, genes were sorted by log fold change and the top 50 AR-specific, GR-specific and shared target genes between AR and GR with highest fold change were plotted using the pheatmap and ggplot2 packages. For the non-regulated gene category, RNA-seq heatmaps of 50 randomly selected genes were plotted. Similarly, heatmaps for GR-specific RIME interaction partners were generated.

#### Reviewers access to new datasets submitted to EBI ArrayExpress

RNA-seq in U2OS-AR (24h R1881/dmso): E-MTAB-9622

Username: Reviewer_E-MTAB-9622

Password: QK36V9gk

FAIRE-STARR-seq in U2OS-AR and U2OS-GR (14h R1881/dmso or 14h dex/etoh): E-MTAB-9614

Username: Reviewer_E-MTAB-9614

Password: 79adRziv

ATAC-seq in U2OS-AR (4h R1881/dmso): E-MTAB-9606

Username: Reviewer_E-MTAB-9606

Password: osLskfby

AR ChIP-seq in U2OS-AR (4h R1881, 2 replicates) + GR ChIP-seq in U2OS-GR (1.5h dex, 1 replicate): E-MTAB-9616

Username: Reviewer_E-MTAB-9616

Password: hnyuFut1

All histone modifications (K27ac, K4me1, K9me3, K27me3) in U2OS-AR and U2OS-GR (hormone+vehicle each, inputs): E-MTAB-9617

Username: Reviewer_E-MTAB-9617

Password: vvesoyUQ

## Results

### Transcriptional Regulation and Genomic Binding by GR and AR

To study how TF paralogs can diverge functionally despite having nearly identical DNA binding domains, we used the same parental cell line to generate cells that either express GR (Rogatsky et al. 1997) or AR (Fig. 1A, Fig. S1). Characterization of these cell lines, using whole-cell [^3^H] steroid binding assays, showed that AR levels were about 3 times lower than for GR (Fig. 1B). Further, robust regulation of the *FKBP5* and other target genes was observed 4 hours after hormone treatment of the U2OS-GR line. In contrast, regulation of FKBP5 and other genes required a markedly longer hormone treatment for the U2OS-AR line (Fig. 1C). Given the slower kinetics of gene regulation for AR, we decided to generate and compare RNA-seq data for U2OS-GR cells treated for 4 hours and U2OS-AR cells treated for 24 hours. For upregulated genes, we identified three classes of genes: GR-specific genes (590) AR-specific genes (177) and genes regulated by both AR and GR (187 genes), (Fig 1D, Fig. S1). The number of repressed genes was much smaller for AR (17) than for GR (230) with little overlap (3 genes) between the two gene sets.

**Fig. 1.**
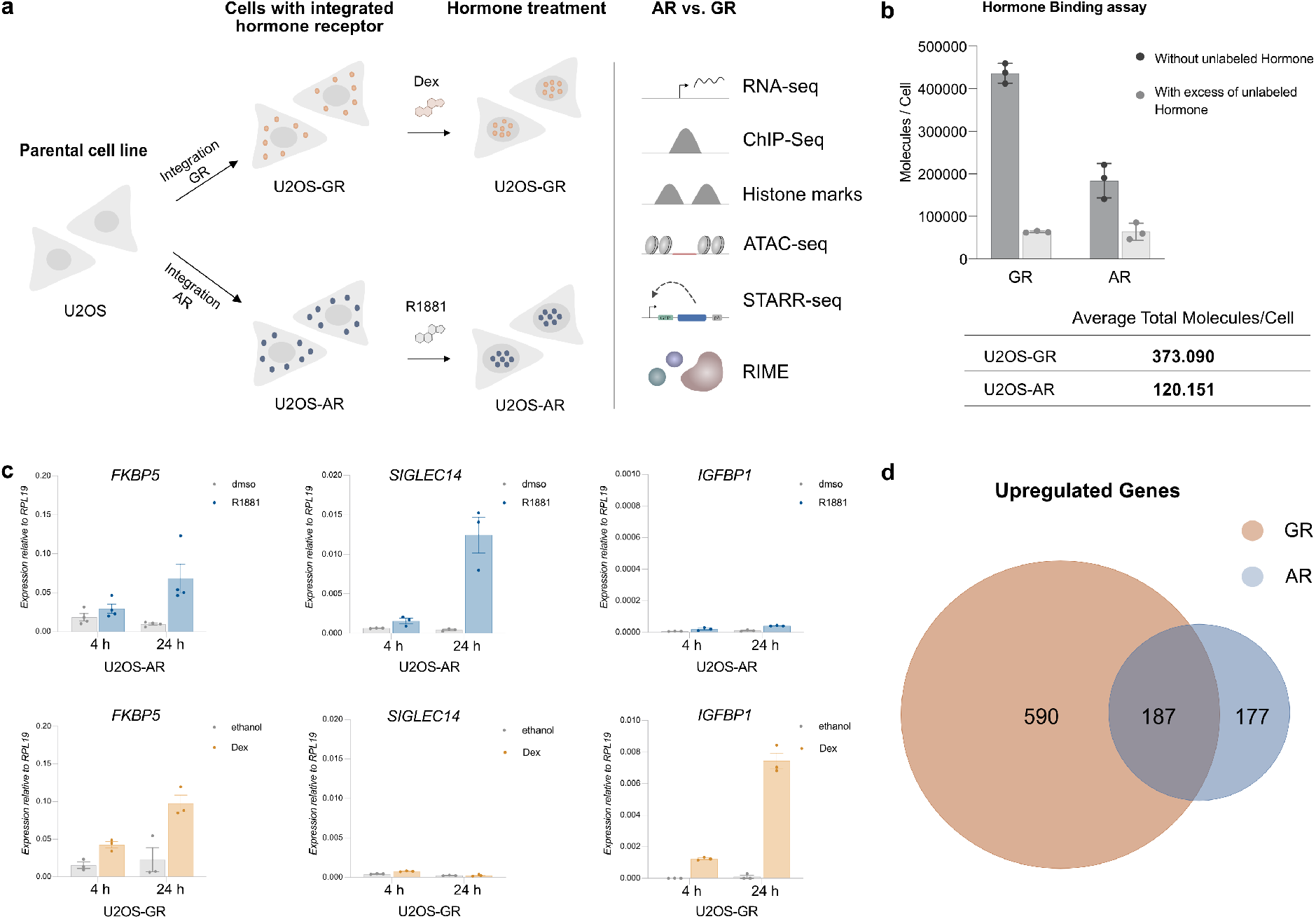
Establishment of AR cell line & comparison gene regulation AR vs GR. (a) Experimental design. Cell lines expressing either AR or GR were derived from the parental U2OS human bone osteosarcoma cell line and used for a variety of experiments as shown. (b) The number of bound ligand molecules per cell was determined for the U2OS-AR and U2OS-GR cell lines by treating cells with either 100 nM [3H-]-R1881 (for AR) or 100 nM [3H-]-Dex (for GR) in the presence or absence of a 10 μM excess of the corresponding unlabeled ligand. The number of total molecules per cell was calculated by subtracting the average number of molecules per cell with excess of unlabeled hormone from the average number of molecules per cell without excess of unlabeled hormone. The average of three independent replicates ±SEM is shown. Here and elsewhere, dots depict values for each individual experiment. (c) Relative mRNA levels of *FKBP5*, *SIGLEC14* and *IGFBP1* was quantified by qPCR for U2OS cells stably expressing either (top) AR or (bottom) GR. U2OS-AR cells were treated for 4h or 24h with dmso as vehicle control or 5 nM R1881. U2OS-GR cells were treated with ethanol as vehicle control or 1 μM Dex for 4h or 24h. Average gene expression ±SEM is shown (n ≥ 3). (d) Venn diagram showing the overlap in genes upregulated by AR and GR based on RNA-seq data. U2OS-AR cells were treated for 24h with 5 nM R1881; U2OS-GR cells were treated for 4h with 1 μM Dex. Genes were designated as significantly upregulated when the adjusted p-value < 0.05 and |log2(fold change) | > 1.5.

Differential patterns of genomic occupancy for AR and GR likely play a role in directing receptor-specific gene regulation (Sahu et al. 2013; Zhang et al. 2018). To compare the cistromes between AR and GR, we generated and analyzed ChIP-seq data for both hormone receptors. We called peaks and created three peak categories: shared peaks (peak called in each of the two replicates for both AR and GR), AR-specific peaks (peak called in both AR replicates, but not for GR) and GR-specific peaks (peak called in both GR replicates, but not in AR). Consistent with what we observed in terms of gene regulation, a substantial fraction of AR binding peaks overlaps with GR-occupied loci (Fig. 2A). In addition, we find a category of AR-specific peaks and a large number of binding sites that are occupied in a GR-specific manner (Fig. 2A,B).

**Fig. 2.**
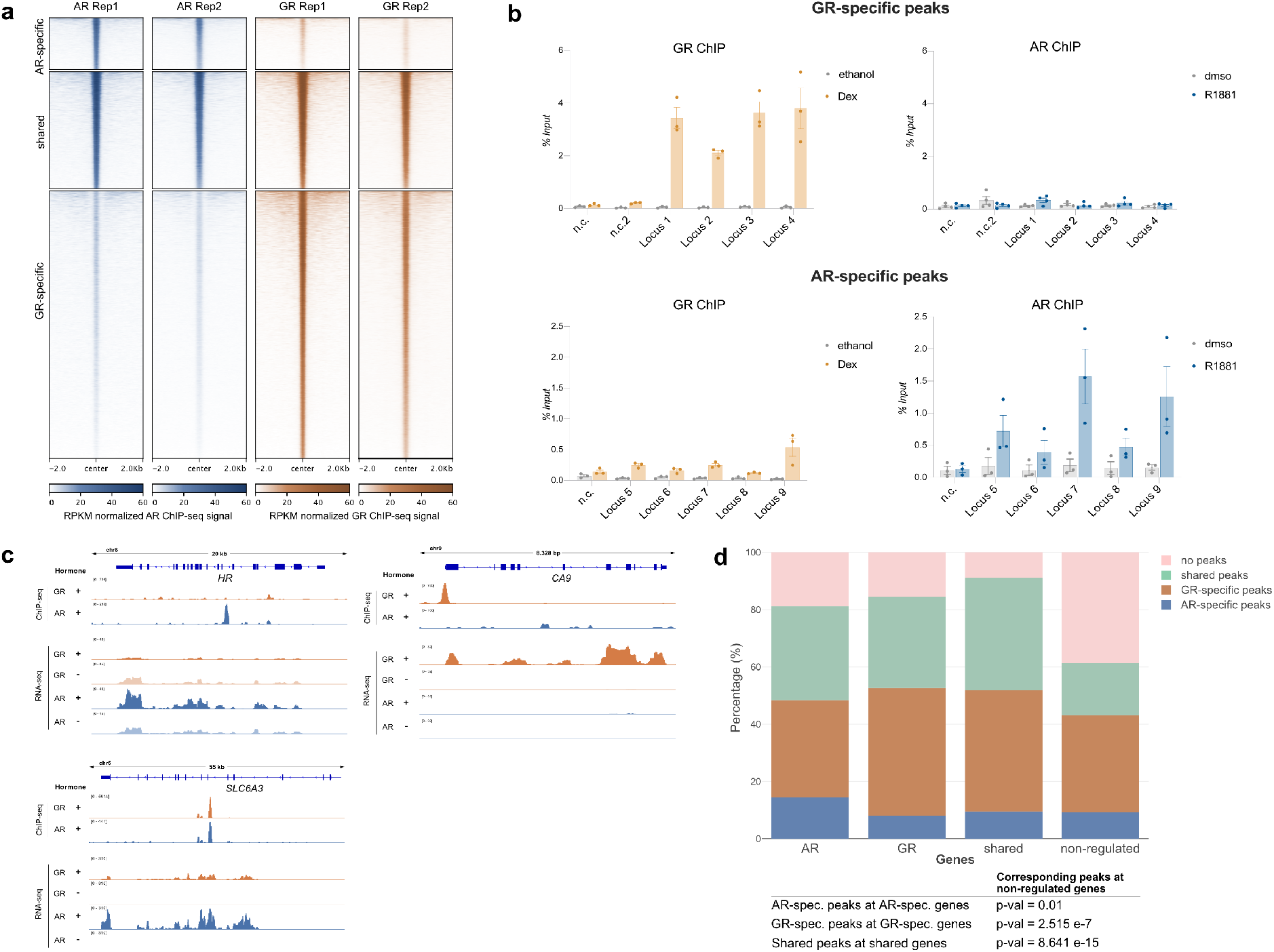
Comparison genome-wide GR binding. (a) Heatmap visualization of AR and GR ChIP-seq read coverage (RPKM normalized) at shared and receptor-specific binding sites (+/− 2 kb around peak center). U2OS-AR cells were treated with R1881 (5 nM, 4h) and U2OS-GR cells with Dex (1 μM, 1.5h). (b) Validation of GR-specific peaks (top) or AR-specific peaks (bottom) by ChIP-qPCR. U2OS-GR cells were treated with 1 μM Dex or ethanol as a vehicle control for 1.5h. U2OS-AR cells with 5 nM R1881 or dmso for 4h. Averages ± SEM are shown (n ≥ 3). (c) Examples of AR-specific (*HR*), GR-specific (*CA9*) and shared (*SLC6A3*) genes are shown as genome browser screenshots depicting the ChIP-seq data (top) and RNA-seq data (bottom). For ChIP-seq experiments, U2OS-GR cells were treated with ethanol or 1 μM Dex for 1.5h. U2OS-AR cells with dmso or 5 nM R1881 for 4h. One representative ChIP-seq track is shown from two biological replicates. For the RNA-seq analysis, U2OS-GR cells were treated with ethanol or 1 μM Dex for 4h. U2OS-AR cells with dmso or 5 nM R1881 for 24h. Merged RNA-seq track from three biological replicates is shown. (d) Stacked bar graphs showing the distributions of different categories of peaks (shared, GR-specific, AR-specific and no peaks) for each category of regulated genes (AR-specific, GR-specific, shared and non-regulated). p-values were calculated using a Fisher’s exact test.

Next, we assessed whether differential occupancy contributes to the receptor-specific transcriptional regulation we observed. Given the small number of repressed genes for AR and the ambiguous link between GR binding and transcriptional repression (Sasse et al. 2019), we decided to focus our analysis on upregulated genes. Genes were categorized as either non-regulated by either AR or GR, shared between AR and GR, GR-specific or AR-specific (Fig. S1). For each gene within a category, we scanned a 60 kb window centered on the transcriptional start site for the presence of either a peak shared by AR and GR, a GR-specific peak, an AR-specific peak or the absence of a peak.

As expected, we found that a larger fraction of non-regulated genes contains no peaks in this window than genes in the other categories, indicating that nearby receptor binding correlates with gene activation (Fig. 2D). Furthermore, AR-specific binding is enriched near AR-specific genes whereas GR-specific binding is enriched near GR-specific genes as well as shared target genes (Fig. 2C,D). Together, these data indicate that receptor-specific binding is a driver of receptor-specific gene activation. However, for the majority of genes that are activated specifically by either AR or GR, we do not observe receptor-specific binding suggesting that receptor-specific regulation might also be governed by events downstream of binding.

### Role of chromatin in shaping genome-wide receptor binding

To investigate the role of chromatin accessibility in shaping the genome-wide binding of AR and GR, we performed ATAC-seq (Buenrostro et al. 2013) and ChIP-seq experiments for a panel of histone modifications for vehicle-treated cells to capture the chromatin landscape the receptors encounter when activated by hormone. Next, we intersected the ATAC-seq data with the three peak categories (shared; AR-specific; GR-specific binding). The most striking difference between the peak categories is that GR-specific peaks are, on average, markedly less accessible prior to hormone treatment than either AR-specific or shared peaks (Fig. 3A). We also performed ChIP-seq experiments for a panel of histone modifications that are associated with either closed or open chromatin (ENCODE Project Consortium 2012). Consistent with preferential GR-specific binding at relatively inaccessible loci (Fig. 3A), the levels of histone modifications associated with closed chromatin (H3K27me3 and H3K9me3) are higher for the GR-specific peaks than for either shared or AR-specific peaks (Fig. S2A,B).

**Fig. 3.**
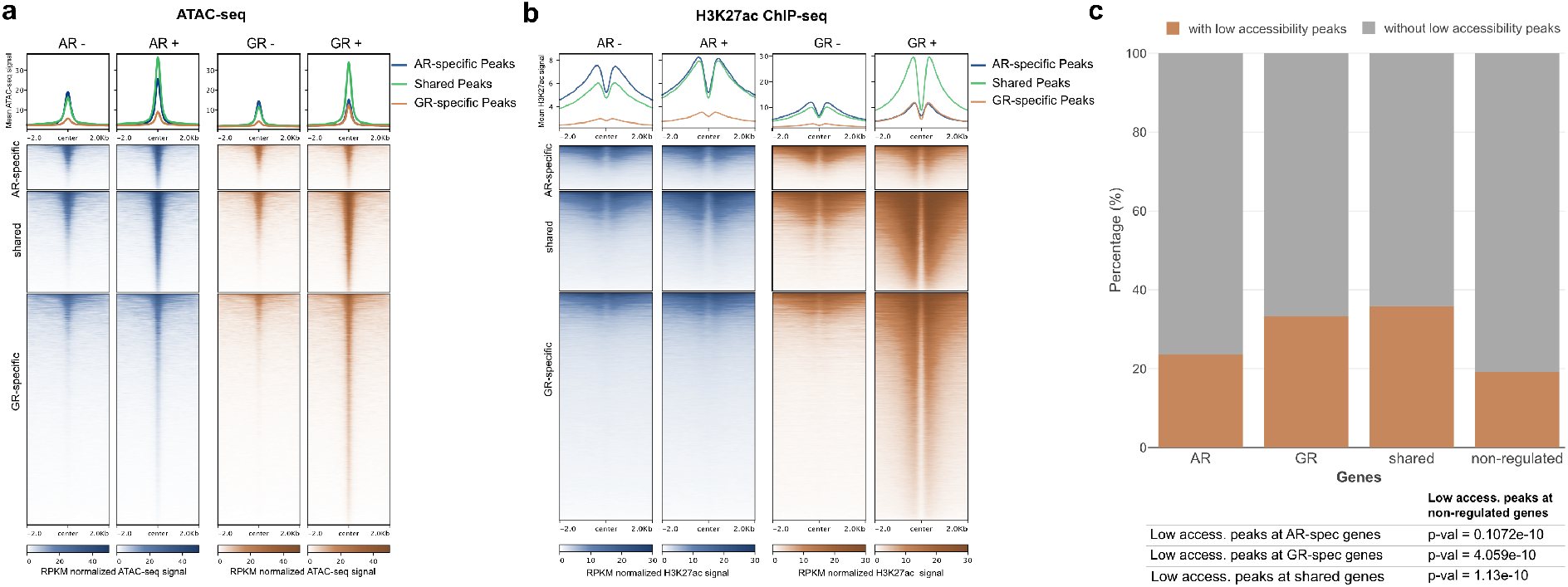
Causes & consequences of receptor binding. (a,b) Heatmap visualization and (top) mean signal plot of (a) ATAC-seq or (b) H3K27ac ChIP-seq read coverage (RPKM normalized) at shared and receptor-specific binding sites (+/− 2 kb around peak center). U2OS-AR cells were treated with R1881 (5 nM, 4h: AR +)) or vehicle (AR -); U2OS-GR cells were treated with Dex (1 μM, 1.5h: GR +) or vehicle (GR -). (c) Stacked bar graphs showing the percentage of genes with a “low accessibility GR peak” in a 60kb window centered on the TSS of each gene for each category of regulated genes (AR-specific, GR-specific, shared and non-regulated). P-values were calculated using a Fisher’s exact test.

Conversely, the levels of histone marks associated with open chromatin (H3K4me1, H3K27ac, (ENCODE Project Consortium 2012)) are lower for the GR-specific peaks (Fig 3B, Fig. S2C). Together, these results argue that a different propensity to bind relatively inaccessible chromatin plays a role in directing receptor-specific binding.

In our set-up, GR levels are about three times higher than AR levels (Fig. 1B). Therefore, we wanted to test if GR-specific binding at relatively inaccessible chromatin is a simple consequence of higher GR levels, or an intrinsic property that distinguishes GR from AR. To test if GR can still bind when reduced levels of hormone-occupied receptor are present, we assayed GR occupancy at hormone concentrations below the reported *KD*s of GR for dexamethasone (∼3-5 nM, (Koubovec et al. 2005; Vedder et al. 1993)). We first confirmed GR-specific binding at 4 low-accessibility loci when a saturating amount of hormone was used (Fig. S3A). At lower hormone concentrations (0.5 nM and 1 nM), GR occupancy was reduced but still detectable at each of the loci examined (Fig. S3A). Previous studies have shown that GR binding induces increased chromatin accessibility (John et al. 2008). Consistent with GR-specific binding, we observed an increase in chromatin accessibility upon hormone treatment at these 4 low-accessibility loci for GR but not for AR (Fig. S3B). The increase was smaller, but still observable, at sub-saturating hormone concentrations (1 nM) arguing that binding and opening is not a simple consequence of higher receptor levels for GR than for AR.

Next, we investigated if GR binding at regions of low chromatin accessibility might contribute to GR-dependent gene regulation. Therefore, we filtered GR-specific peaks for those mapping to relatively inaccessible chromatin (“Pioneering GR-peaks”) and intersected them with the different categories of regulated genes. In line with a role in regulating gene expression, we found that these low-accessibility GR peaks are enriched near genes regulated by GR (Fig. 3C). Together, these results argue that a different propensity to bind inaccessible chromatin plays a role in directing receptor-specific binding and gene regulation.

### Role of sequence in shaping genome-wide receptor binding

Differences in DNA-binding specificity between related TFs can induce differential binding and gene regulation (Rohs et al. 2010). To study the role of sequence composition in directing AR and GR to different genomic loci, we used AME (Analysis of Motif Enrichment, (McLeay and Bailey 2010)) to scan the clustered JASPAR CORE vertebrates motif collection (Castro-Mondragon et al. 2017) supplemented with a direct repeat AR/GR consensus motif (DR3) which is reported to be AR-specific (Schoenmakers et al. 2000). Similar to a previous study comparing the motif composition of AR- and GR-specific sites (Sahu et al. 2013), we found that enriched sequence motifs largely overlap between AR- and GR-specific sites with some differences (Fig. S4C). For example, the DR3 motif was more enriched for AR-specific binding sites whereas the canonical inverted AR/GR consensus motif was more enriched for GR-specific sites. The most striking difference was for the AP-1 motif which was the most enriched motif for AR-specific sites with little enrichment for GR-specific sites. Given the role of AP-1 in maintaining open chromatin (Biddie et al. 2011), the lack of motif enrichment for GR-specific sites could reflect preferential binding at inaccessible chromatin. To remove chromatin accessibility as a potential confounding factor, we repeated the motif analysis using GR-specific sites at open chromatin regions which show a similar level of ATAC-seq signal as the AR-specific binding sites (Fig. 4A).

**Fig. 4.**
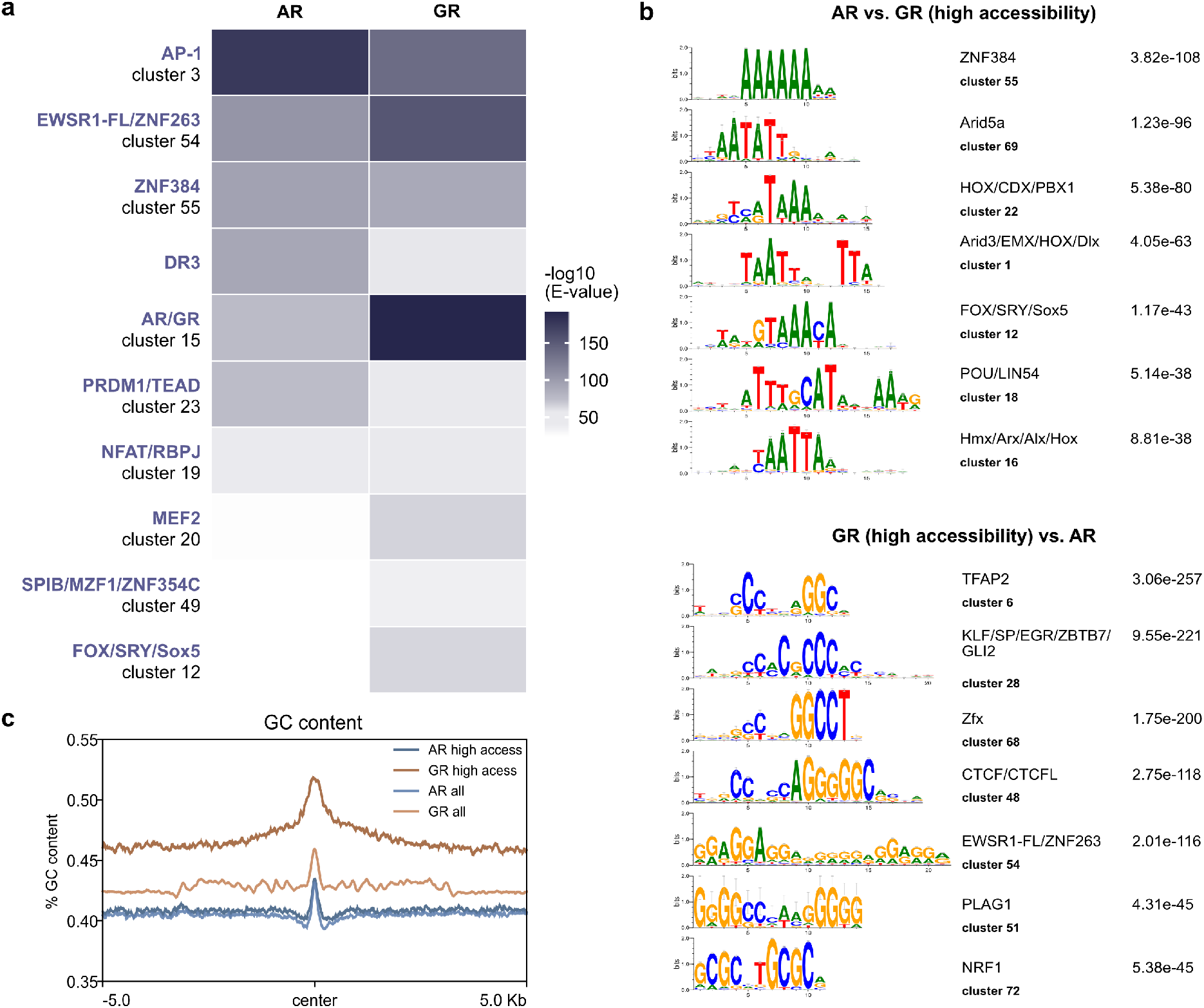
Role of sequence in directing receptor-specific binding. (a) Heatmap visualization of enriched motif clusters at all AR-specific and high-accessibility GR-specific peaks (+/− 250 pb around the peak center). Shuffled input sequences were used as background for the motif enrichment analysis. Motifs were included if the E value was <10-30 for either AR or GR. (b) Top: Top 7 enriched differential motif clusters and corresponding E value at all AR-specific peaks when compared to an equal number of high-accessibility GR-specific peaks (+/− 250 pb around the peak center). Bottom: Top 7 enriched differential motif clusters at high accessibility GR-specific peaks when compared to all AR-specific peaks (c) Mean GC content at all AR-specific peaks (AR all), all GR-specific peaks (GR all) and peaks in regions of accessible chromatin (AR high access or GR high access) (+/− 5 kb around the peak center). Based on Mann-Whitney-U test, GC content is significantly higher for GR-specific peaks (p-value < 2.2e-16).

Analysis of selective receptor binding in regions with similar chromatin accessibility showed motif hits that resembled the results for all binding sites except, for example, that the GR-specific sites in open chromatin were also enriched for AP-1 (Fig. 4A).

Next we repeated the AME analysis however this time using AR-specific peaks as background for GR-specific peaks (either all peaks or only GR peaks in accessible chromatin) and vise-versa to identify motifs that are selectively enriched. Interestingly, the top AR-specific motifs were mostly AT-rich whereas the top motifs for GR-specific peaks were often high in GC content suggesting that the sequence composition of selectively occupied regions is different (Fig. 4B). Accordingly, when we scanned the GC content of AR and GR-specific genomic regions, we found a higher GC content for GR occupied regions than for regions occupied by AR (Fig. 4C). This difference was most pronounced when we compared receptor-specific peaks in regions of accessible chromatin (Fig. 4C). This finding is in line with a recent study showing that AR binding is distinguished from GR by a preference for poly(A) sequences directly flanking the consensus binding site (Zhang et al. 2018). Surprisingly, in contrast to the *in vitro* study we find that the difference extends far beyond the core motif and its direct flanks (>10kb) arguing that the sequence composition of a large region surrounding the binding site might play a role in directing receptor-selective recruitment.

### Receptor-specific consequences of DNA binding

To compare the events that occur downstream of AR and GR binding, we assayed the effect of hormone treatment on several chromatin features. First, we compared changes in chromatin accessibility by analyzing ATAC-seq signal after hormone treatment. In agreement with published data (John et al. 2011; Tewari et al. 2012), both AR and GR induce an increase in chromatin accessibility at occupied loci (Fig. 3A). For GR, accessibility increased at shared and GR-specific sites but not at AR-specific regions. For AR, the increase in accessibility was most pronounced at shared and AR-specific peaks however, a slight increase was also observable for the GR-specific sites indicating that some AR binding also occurs at some of these loci.

Next, we analyzed the effects of hormone treatment on histone modifications. Specifically, we analyzed H3K27ac as a marker for active enhancers and indicator of enhancer activity as well as H3K4me1 as a marker of active and primed enhancers (Creyghton et al. 2010; Rada-Iglesias et al. 2011). Consistent with increased chromatin accessibility at occupied loci, we found that GR activation by hormone, and to a lesser degree for AR, induced nucleosome shifts for H3K4me1 and H3K27ac resulting in reduced signal at the center of the receptor-occupied locus relative to the regions directly flanking it (Fig. 3B, Fig. S2C). In line with expectation, GR activation resulted in a marked increase in H3K27ac levels at GR-occupied loci (Fig. 3B). In contrast, H3K27ac levels barely changed in response to AR activation at AR-occupied loci (Fig. 3B). To test if the receptor-specific changes in H3K27ac levels are restricted to the time-point assayed, we tested several loci occupied by both AR and GR by ChIP-qPCR. Specifically, we chose 4 loci located near genes that are regulated by both receptors (Fig. S5A) and assayed H3K27ac levels after both 4h and 24h of hormone treatment. Consistent with our genome-wide analysis, a marked increase in H3K27ac levels was observed 4 hours after GR activation with even higher levels after 24h (Fig. S5B). For AR, H3K27ac levels did not change after 4h of hormone treatment whereas relatively small increases were observable after 24h. Together, these results argue that both AR and GR induce chromatin remodeling and increased chromatin accessibility upon genomic binding. However, the consequences downstream of AR and GR binding diverge when H3K27ac levels are examined with robust and rapid increases for GR whereas AR activation only induces modest changes in H3K27ac levels that occur with slower kinetics.

Although several studies indicate that H3K27ac levels are indicative of enhancer activity (Creyghton et al. 2010; Rada-Iglesias et al. 2011) some studies argue that H3K27 acetylation is dispensable for enhancer activity (Zhang et al. 2020). For a quantitative comparison of the ability of receptor-occupied regions to enhance transcription, we performed STARR-seq (Self-Transcribing Active Regulatory Region sequencing) for both AR and GR with two modifications (Fig. 5A). First, to limit the number of putative enhancers we focused on genomic regions isolated by FAIRE (Formaldehyde Assisted Isolation of Regulatory Elements) from dexamethasone-treated U2OS-GR cells to include regions that gain accessibility upon GR activation. Second, we added unique molecular identifiers (UMIs) during the reverse transcription stage to facilitate quantitative measurements of enhancer activity for each fragment. To quantify enhancer activity, we transfected U2OS-AR and U2OS-GR cells with the FAIRE-reporter library and assayed regulatory activity for vehicle-treated cells and for cells treated overnight with the corresponding hormone. Intersection of the STARR-seq data with the different groups of receptor binding sites, showed that enhancer activity increased upon dexamethasone treatment for GR-occupied regions whereas the enhancer activity of the group of AR-specific peaks did not change (Fig. 5B). For AR, we observed increased enhancer activity for the shared and AR-specific sites upon R1881 treatment. In addition, AR activates enhancers of the GR-specific group of binding sites indicating that AR can bind these regions when they are taken out of their endogenous chromatinized genomic context.

**Fig. 5.**
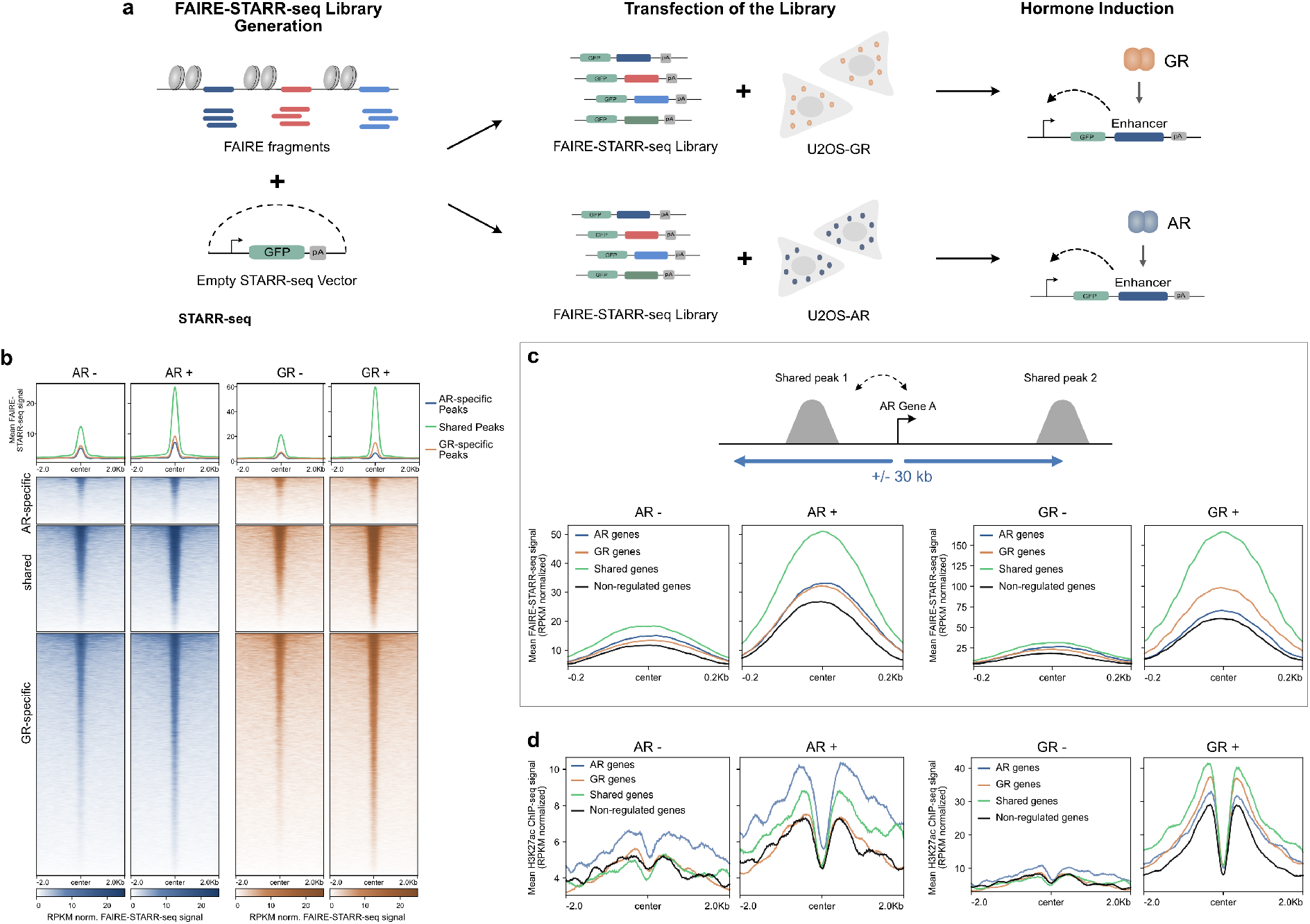
Comparing intrinsic enhancer activity between AR and GR. (a) FAIRE STARR-seq experimental set-up (see Methods for details). (b) Heatmap visualization and mean signal plot of FAIRE-STARR-seq read coverage (RPKM normalized) at shared and receptor-specific binding sites (+/− 2 kb around the peak center). U2OS-AR cells were treated with R1881 (5 nM, 14h: AR +) or vehicle (AR -). U2OS-GR cells with Dex (1 μM, 14h: GR +) or vehicle (GR -). (c) Top: Cartoon showing how shared peaks were assigned to each category of regulated genes (AR-specific, GR-specific, shared or non-regulate). Bottom: Mean signal plot of FAIRE-STARR-seq read coverage (RPKM normalized) at shared sites (+/− 250 bp around the peak center) near the different categories of regulated genes. AR -: STARR-seq coverage for vehicle treated U2OS-AR cells. AR +: same for R1881 treated cells; GR -: STARR-seq coverage for vehicle treated U2OS-GR cells. GR +: same for Dex treated cells (d) Same as for (c) except that the mean H3K27ac ChIP-seq read coverage (RPKM normalized) at shared sites is shown (+/− 2 kb around the peak center).

Quantitatively, at the time point examined, the overall level of activation is higher for GR than for AR. However, despite the very modest induction of H3K27 acetylation for AR, enhancer activation is also observed for AR. Moreover, an exemplary enhancer near the AR-specific *AQP3* gene shows activation by AR but not by GR (Fig. 7A, Fig. S5E) in the STARR-seq assay whereas H3K27ac changes upon binding are much more pronounced for GR indicating that enhancer activation and H3K27 acetylation can be uncoupled.

Notably, both regulated- and non-regulated genes harbor proximal receptor binding sites (Fig. 2D) arguing that events downstream of binding play a role in specifying if a nearby gene is regulated or not. To test if the enhancer activity of shared peaks correlates with the regulation of nearby genes, we compared shared peaks that are located near genes that are either non-regulated, shared targets of AR and GR or receptor-specifically regulated (Fig. 5C). Consistent with a role of of enhancer activity in directing changes in gene expression of nearby genes, we find that for both AR and GR the STARR-seq activity of shared peaks near regulated genes is higher than for shared peaks that are located near non-regulated genes (Fig. 5C). Furthermore, for GR enhancer activity after hormone treatment is highest for shared peaks near shared and GR-specific genes with lower activity at shared peaks near AR-specific genes and non-regulated genes (Fig. 5C). Similarly, enhancer activity for AR is highest for shared peaks near shared genes followed by AR-specific genes, GR-specific and finally non-regulated genes. Further arguing for a role for events downstream of binding in directing specificity, we find that H3K27ac levels (Fig. 5D) and ATAC-seq signal (Fig. S5C) at shared peaks after hormone treatment correlate with receptor-specific regulation. The ChIP-seq signal for AR and GR at shared peaks is also a bit higher at peaks near regulated genes, indicating that receptor levels likely contribute to the differences in enhancer activity (Fig. S5D).

Together, these results show that enhancer activity of receptor-occupied regions correlates with transcriptional activation of genes. Comparison between AR and GR shows that activity is highest near shared genes. For GR, peaks near GR-specific genes are more active than those located near AR-specific genes. This order is reversed for AR, arguing that receptor-specific enhancer activity at shared binding sites plays a role in directing receptor-specific regulation of nearby genes.

### Shared binding sites contribute to receptor-specific target gene regulation

The presence of shared binding sites near receptor-specific target genes prompted us to explore experimentally if shared binding sites contribute to the regulation of the nearby gene. Specifically, we picked the GR-specific target gene *GILZ* and the AR-specific gene *AQP3* and confirmed the receptor-specific regulation by qPCR (Fig. 6A). For *GILZ*, each of the GR-occupied peaks in an approximately 100 kb window centered on the regulated promoters is also occupied by AR (Fig. 6B). We previously showed that simultaneous deletion of the promoter-proximal peaks and a region downstream of *GILZ* containing multiple peaks resulted in a virtual loss of GR-dependent regulation (Thormann et al. 2018). Re-examination of the clonal lines confirmed that the proximal enhancer and distal GR binding sites in the downstream region are required for GR-dependent regulation of *GILZ* but not of *FKBP5*, a GR target gene located on another chromosome (Fig. 6C). Thus, despite occupancy for both AR and GR at each of the peaks required for GR-dependent regulation, AR fails to regulate *GILZ*. To assess whether differential enhancer activity could explain receptor-specific transcriptional regulation, we compared the STARR-seq signal at shared peaks for hormone-treated cells (Fig. S6A). For several shared peaks of the *GILZ* gene, the STARR-seq activity upon hormone treatment appears higher for GR than for AR indicating that the level of enhancer activity at shared binding sites might play a role in directing GR-specific regulation.

**Fig. 6.**
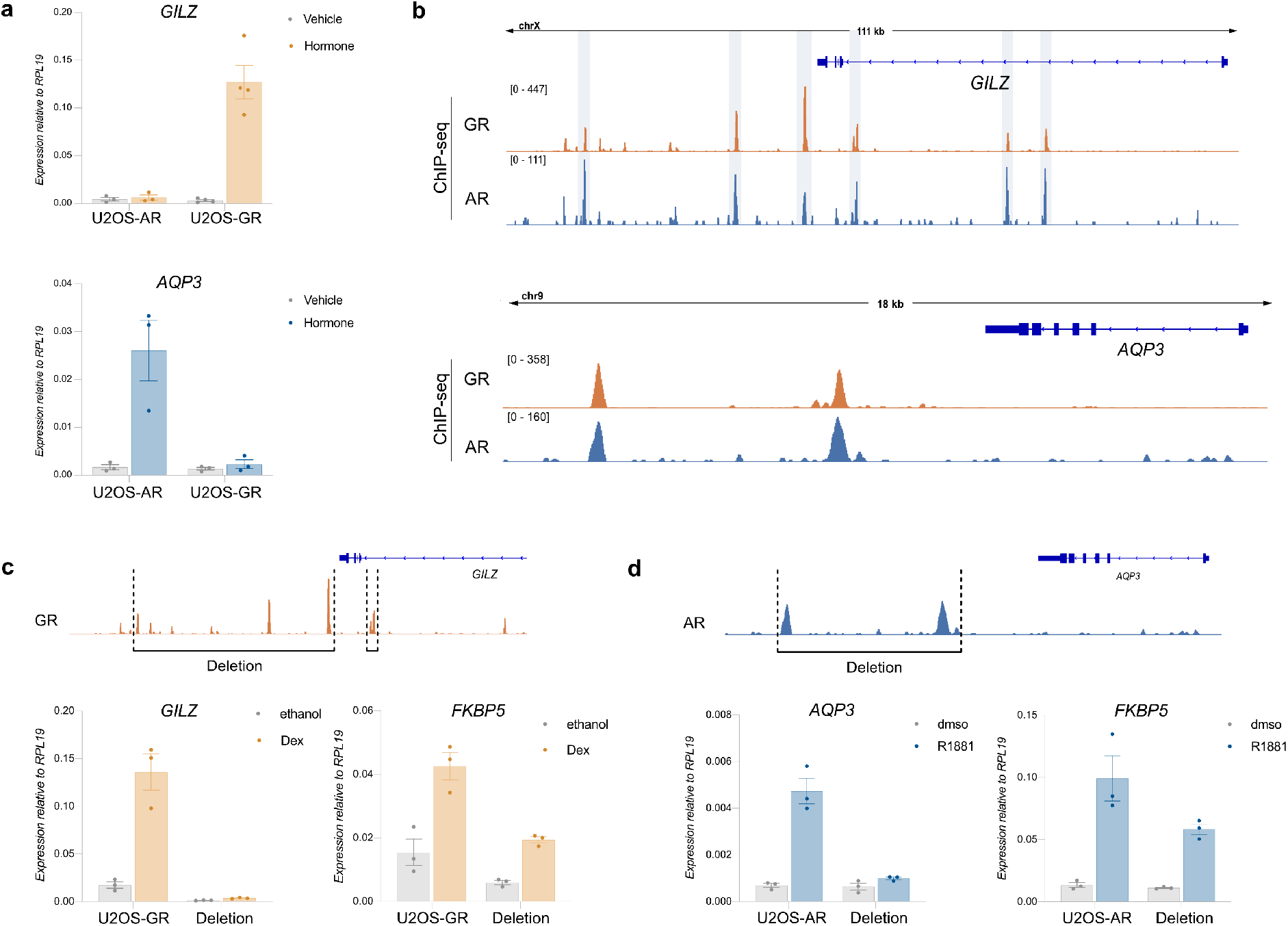
Shared binding sites direct receptor-specific gene regulation. (a) Relative mRNA levels of *GILZ* and *AQP3* was quantified by qPCR as described for Fig. 1C. Averages for cells treated for 24h ±SEM are shown (n ≥ 3). (b) ChIP-seq read coverage (RPKM normalized) for GR and AR for (top) the GR-specific target gene *GILZ* and (bottom) the AR-specific target gene *AQP3*. Cells were treated as specified for Fig. 2c. (c) Top: GR ChIP-read coverage (RPKM normalized) highlighting the regions that were deleted for the *GILZ* deletion clonal lines in U2OS-GR cells. Bottom: Relative mRNA levels as determined by qPCR for the *GILZ* and *FKBP5* genes are shown for wt U2OS-GR and the *GILZ* deletion clonal lines. Cells were treated for 4h with 1 μM Dex or ethanol as vehicle control (a). Averages ±SEM are shown. (d) Same as (c) except for *AQP3* deletion clonal lines in U2OS-AR cells. Cells were treated for 24h with 5 nM R1881 or dmso as vehicle control.

For the AR-specific *AQP3* gene, two prominent shared peaks occupied by both AR and GR are located in a region approximately 5-10 kb downstream of the transcriptional start site (Fig. 6D). To test if these peaks contribute to AR-dependent regulation, we used CRISPR/Cas9 (Mali et al. 2013) and a pair of sgRNAs to remove a ∼4 kb genomic fragment containing both peaks (Fig. 6D, Fig. S6B). Analysis of the resulting clonal lines showed that AR no longer regulates the *AQP3* gene when these peaks are deleted (Fig. 5D) whereas the AR target gene *FKBP5*, which is located on another chromosome, is still regulated by AR. Interestingly, the peak closest to the *AQP3* gene (“*AQP3* enhancer”) shows an increase in STARR-seq signal in U2OS-AR cells upon hormone treatment whereas no such increase is observed for GR (Fig. 7A). To characterize the *AQP3* enhancer, we constructed a reporter containing a 554 bp region centered on the peak.

**Fig. 7.**
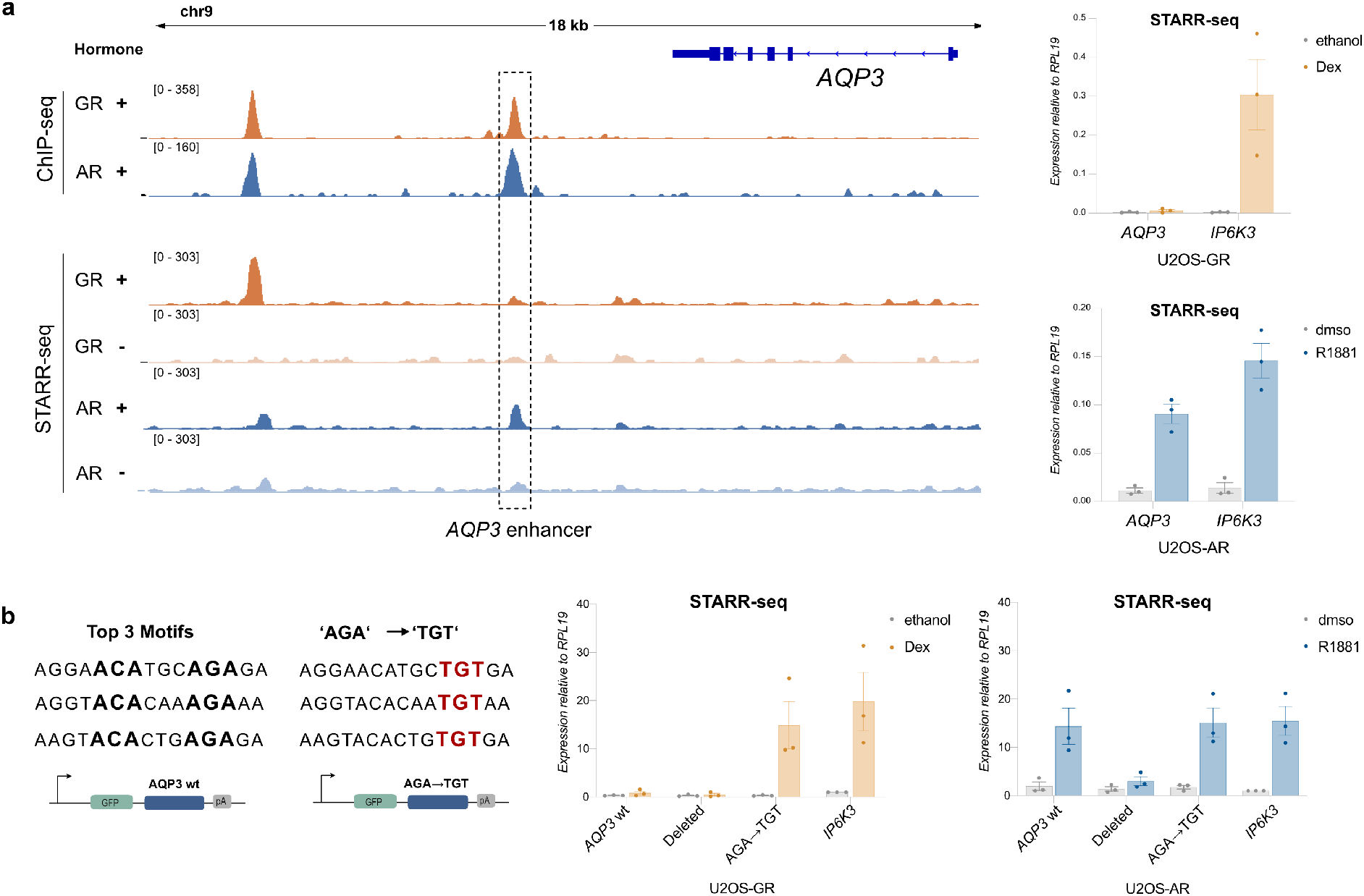
Mutational analysis of the AQP3 enhancer. (a) Left: ChIP-seq and FAIRE STARR-seq read coverage (RPKM normalized) for GR and AR at the *AQP3* locus. The enhancer with AR-specific activity is highlighted. Cells for ChIP-seq were treated as for Fig. 2A. Treatment for FAIRE STARR-seq as described for Fig. 5B. Right: Transcriptional activity of STARR-seq reporters containing either the *APQ3* enhancer or an enhancer near the *IP6K3* gene that is activated by both AR and GR. Relative mRNA levels ± S.E.M. are shown for cells treated overnight with either vehicle or with 5 nM R1881 (U2OS-AR cells) or 1 μM Dex (U2OS-GR cells). Averages ± SEM are shown (n = 3). (b) Left: Top 3 AR motif (JASPAR MA0007.1-3) matches of the *AQP3* enhancer region. Positions highlighted in bold were changed to ATA to delete each of the three motif matches (Deleted). AGA sequence was mutated to TGT to create motifs resembling the canonical GR consensus motif (AGA -> TGT). Right: Transcriptional activity of STARR-seq reporters as indicated for GR and AR. Cells were treated as for (a). Averages ± SEM are shown (n = 3).

Testing the activity of this reporter confirmed that the *AQP3* enhancer is activated in an AR-specific manner whereas an enhancer near the *IP6K3* gene was regulated by both AR and GR (Fig. 7A). To test the influence of sequence in directing the observed receptor-specific regulation of the *AQP3* enhancer, we scanned the sequence for occurrences of the AR consensus motif and deleted the top 3 matches (Jaspar MA0007.2, Fig. 7B). Each of these three sites contains one well-defined half-site with a more degenerate second half site. Simultaneous deletion of all 3 motif matches by mutating key positions resulted in a loss of hormone-dependent activation for AR showing that these binding sites are required for regulation (Fig. 7B). Next, we mutated each of the top 3 matches to resemble the GR consensus motif with a well-defined second half-site (Fig. 7B, AGA -> TGT). This mutated *AQP3* reporter could now be activated by both AR and GR, indicating a role of these sequences in directing AR-specific regulation of the reporter. Together, these results argue that bound regions that are shared between AR and GR play a role in directing receptor-specific regulation.

### RIME uncovers different interactomes for AR and GR

AR and GR have very similar DNA binding domains whereas the rest of the protein, especially the N-terminal domain, is much less conserved. As a consequence, AR and GR might interact with different transcriptional co-regulators. This in turn could contribute to receptor-specific regulation, *e.g.* when cofactors that address a rate limiting step in the transcriptional activation of a gene are recruited in a receptor-specific manner. To compare the proteins that interact with GR and AR, we performed RIME (Rapid Immunoprecipitation Mass spectrometry of Endogenous proteins). This method combines formaldehyde fixation of intact cells to stabilize protein complexes with immunoprecipitation of a protein of interest and finally mass spectrometry for protein identification (Mohammed et al. 2013). We performed RIME for hormone-treated U2OS-GR and U2OS-AR cells and identified 105 significantly enriched proteins for GR and 173 for AR (Supplementary files S1_data and S2_data, Fig. 8A). As expected, many of the significantly enriched proteins for GR (59%) were also significantly enriched for AR and enriched gene sets include transcription coactivators, nuclear receptor-coactivators and chromatin remodelers (Fig. 8B). Furthermore, several of the identified proteins are previously validated interactors of GR and AR (*e.g.* ARIDA1, NCOA1, NCOA3, EP300, CREBBP, NCOR1, NCOR2, TRIM28) (DePriest et al. 2016; Petta et al. 2016).

**Fig. 8.**
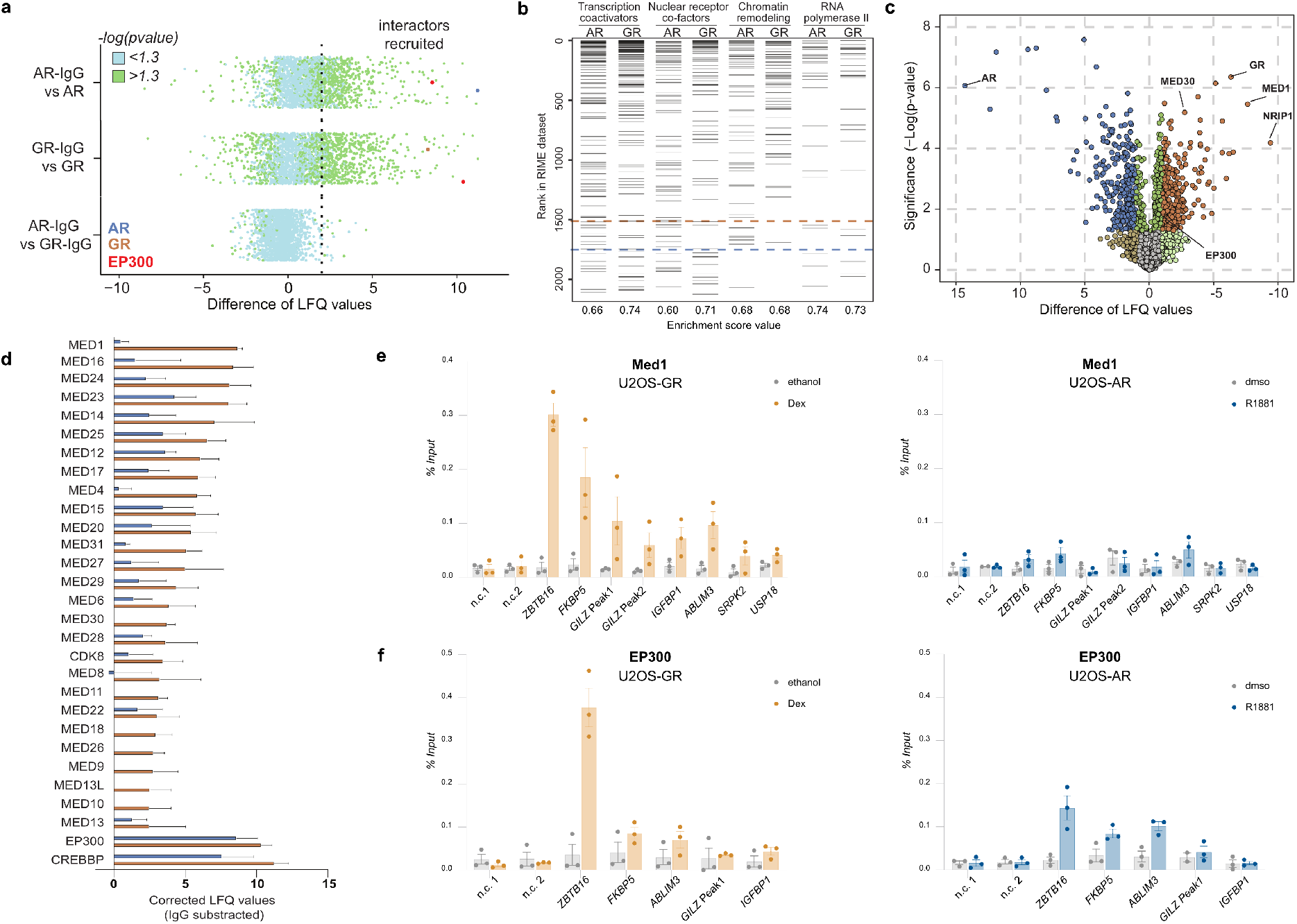
Comparison of AR and GR interactomes. (a) Scatterplot depicting enrichment of GR- and AR-RIME experiments over IgG control. Interactors recruited: ≥ 2 Label-free Quantification (LFQ) enriched over IgG (dotted line) and significant (-log(padj)>1,3; green). n=4. U2OS-GR and U2OS-AR cell lines were treated with either 1μM Dex or 5 nM R1881 for 4h, respectively. (b) Gene Set Enrichment Analysis (GSEA) enrichment ranks for transcription coactivator activity, nuclear receptor binding, chromatin remodeling (M19139), and RNA polymerase transcription factor binding genesets based on GR- and AR-RIME datasets. n=4. (c) Volcano plot depicting differentially enriched interactors for AR and GR. n=4. (d) LFQ enrichment of mediator complex members in GR- and AR-RIME experiments. n=4. (e) Med1 occupancy at loci as indicated was analyzed by ChIP followed by qPCR for cells treated with vehicle control (ethanol for U2OS-GR, dmso for U2OS-AR) or 1 μM Dex, 1.5h (U2OS-GR) or 5 nM R1881, 1.5h (U2OS-AR). Average percentage of input precipitated ± SEM from three independent experiments is shown. (f) same as (e) except that ChIP was for EP300.

Next, we compared the RIME data between AR and GR to identify proteins that interact in a receptor-specific manner (Fig. 8C, supplementary file S3_data). For AR, enriched proteins are linked to RNA splicing and processing whereas for GR, geneset enrichment analysis revealed the mediator complex as the top hit and also included a category of genes linked to acetyltransferase activity (Fig. S7A, supplementary file S3_data). Importantly, enriched proteins in these categories showed comparable expression levels in our RNA-seq data for hormone-treated U2OS-AR and U2OS-GR cells indicating that their enrichment is not a simple consequence of hormone-induced changes in gene expression (Fig. S7B,C). Analysis of the RIME signal indicated that each of the mediator subunits is enriched for GR whereas for AR only a subset is enriched with overall lower signal (Fig. 8D). Notably, GR-specific interactions with mediator subunits have also been reported in other studies (Lempiäinen et al. 2017; Jin et al.2012) indicating that the selectivity of this interaction is not restricted to the cell line we examined. To test if the GR-specific association in our RIME experiments translates into GR-specific recruitment, we performed ChIP experiments for one of the mediator complex members, MED1. Analysis of several GR-occupied loci showed robust MED1 recruitment upon dexamethasone treatment (Fig. 8E). In contrast, no obvious MED1 recruitment was observed for these AR-occupied loci upon R1881 treatment (Fig. 8E).

Given the striking difference in induced H3K27 acetylation between AR and GR (Fig. 3B), we were surprised to see that the enzymes responsible for the majority of H3K27 acetylation, CREBBP and EP300 (Jin et al. 2011), were significantly enriched for both receptors (Fig. 8A,D, Supplementary file S1_data and S2_data). Furthermore, ChIP experiments targeting EP300 showed that it is recruited to several receptor-occupied loci for both AR and GR (Fig. 8F). Thus, the difference in H3K27 acetylation does not appear to be a simple consequence of selective EP300 recruitment by GR but could be due to selective recruitment of other proteins that influence H3K27ac levels or modulation of the activity of EP300/CREBBP.

Taken together, our proteomic data suggests that GR and AR recruit distinct sets of transcriptional co-regulator proteins which may contribute to receptor-specific gene regulation.

## Discussion

Despite their similar modes of action, the physiological consequences of androgen and glucocorticoid signaling diverge. This specificity can be a consequence of tissue specific expression of the receptor (Claessens et al. 2017). However, even in cells that express both AR and GR, the target genes only partially overlap (Sahu et al. 2013; Arora et al. 2013). Here, we show that receptor-specific gene regulation can be facilitated by different mechanisms. First, differences in DNA binding site preference and distinct abilities to bind “closed” chromatin can direct divergent genomic binding patterns and target gene regulation. Second, events downstream of binding facilitate receptor-specific target gene regulation from genomic binding sites that are occupied by both AR and GR. Notably, for the several genes, including *AQP3* and *GILZ*, we find that receptor-specific regulation is observed at more than one hormone concentration and time-point (Fig. S8). None-the-less some of the differences in binding and regulation that we observe might reflect differences in the kinetics between AR and GR.

The differential gene regulation by AR and GR can be explained, in part, by divergence in DNA binding specificity. In line with previous studies showing AR-specific binding *in vitro*, the DR3 motif was more enriched at AR-specific binding sites (Schoenmakers et al. 2000). However, when we examined ChIP-exo for both AR and GR, we found a footprint for both AR and GR indicating that despite the more pronounced enrichment of this motif for AR (Fig. 4A), both receptors might bind to DR3 sequences *in vivo* (Fig. S9). Another difference we observed was that the canonical AR/GR consensus motif was more enriched for GR than for AR suggesting that AR might be able to bind to more degenerate sequences something that has also been described by others (Sahu et al. 2014). In addition, our findings corroborate a recent study showing that AR binding is distinguished from GR by a preference for poly(A) sequences directly flanking the consensus binding site (Zhang et al. 2018). However, whereas this *in vitro* study shows that this preference is restricted to the 3 to 4 base pairs immediately flanking the motif, our *in vivo* binding studies reveal a more global difference in sequence composition between regions that are selectively occupied (Fig. 4C). GR-specific regions are more

GC-rich that AR-specific regions and this difference is even more pronounced when we control for chromatin accessibility which correlates with sequence composition (Fenouil et al. 2012). Moreover, when we analyzed published ChIP-seq data for AR and GR in LNCaP and VCap cells respectively (Sahu et al. 2013), we again found that GR-specific regions have higher GC content than their AR-specific counterpart (Fig. S4D). Interestingly, earlier work also reported global differences in sequence composition that extend far beyond the core binding site when comparing TFs across diverse families (Dror et al. 2015). This suggests that the local environment may help direct TFs from different TF families to distinct genomic loci by yet unknown mechanisms. Our finding that the sequence environment differs between paralogous TFs argues that the sequence environment might also play a role in directing TFs with very similar sequence preferences to distinct binding sites. This might not just apply to AR and GR, but could also play a role in directing paralogous TFs from the homeodomain family to distinct loci given that the global motif environment differs between paralogs (Dror et al. 2015).

Genome-wide TF occupancy is influenced by chromatin environment, with the majority of TF binding occurring at accessible DNA regions (John et al. 2011; Shen et al. 2018; Song et al. 2011; Thurman et al. 2012). Here, we report distinct abilities to bind “closed” chromatin for AR and GR as a potential mechanism that directs receptor-specific binding and gene regulation. Moreover, the shape of the signal for H3K9me3 and H3K27me3 at GR-specific peaks suggests that the sequence motif is embedded in a nucleosomal context (Fig. S2). Thus, similar to so-called pioneering factors (Iwafuchi-Doi and Zaret 2014) and consistent with previous studies, GR appears to interact with nucleosomal motifs (Perlmann and Wrange 1988; Li and Wrange 1993; Johnson et al. 2018). Different abilities to interact with relatively inaccessible chromatin among paralogous TF has also been reported by others. For example, whereas Oct4 can bind “closed” chromatin (Soufi et al. 2012), its homolog from the Pou family of TFs, Brn2, does not (Wapinski et al. 2013). Similarly, the ability to interact with “closed” chromatin differs between members of the Hox family of TFs (De Kumar et al. 2017; Bulajić et al. 2019). One explanation for the paralog-specific binding to “closed” chromatin could be differences in protein expression levels. In support of this hypothesis, a larger fraction of GR peaks maps to “closed” chromatin at saturating hormone concentrations than at hormone concentrations below the *K*_D_ at which only a fraction of GR is hormone-occupied (Reddy et al. 2012). Similarly, we find that GR binding at “closed” loci is weaker at hormone concentrations below the *K*_D_ (Fig. S3). However, even at low hormone concentrations GR binding at these “closed” loci is observable. Accordingly, chromatin accessibility at these loci increases when cells were treated with low hormone concentrations (Fig. S3). Together, these findings indicate that GR binding at “closed” chromatin might not simply be explained by higher expression levels for GR than for AR. A further indication that the lack of AR binding at GR-specific peaks is a consequence of the chromatin context in which they are embedded comes from our STARR-seq experiments. An advantage of the ectopic STARR-seq reporter assay is its ability to assess the enhancer activities of DNA sequences that are silenced endogenously at the chromatin level (Arnold et al. 2013). For GR, increased STARR activity upon hormone treatment is seen for GR-specific peaks, whereas no such increase is observable for AR-specific peaks (Fig 5B) consistent with the receptor-specific occupancy we observed. This is different for AR, which shows an increase in STARR activity upon hormone treatment for AR-specific peaks (Fig. 5B). However, an even stronger activation is seen for the GR-specific peaks arguing that AR is capable of binding and activating transcription from these GR-specific regions when they are removed from their endogenous chromatin context. Thus, a complementary explanation for the paralog-specific binding could be a receptor-specific intrinsic ability to interact with “closed” chromatin. This could be a consequence of receptor-specific interactions with coregulators, *e.g.* chromatin remodelers, that facilitate the binding and or opening of chromatin by GR to induce a feed-forward loop to establish robustly occupancy at these loci. These receptor-specific cofactors might interact with less conserved parts of the receptors, for example the N-terminal part of the receptor which shows little conservation between AR and GR (Kino 2000).

Numerous studies have shown that differential genomic targeting is a mechanism that can generate functional diversification among paralogous TFs. However, paradoxically when comparing the genome-wide binding patterns of paralogous TFs, a large fraction of peaks typically overlaps. For example, when comparing AR and GR, receptor-specific binding only explains receptor-specific regulation for a subset of genes with many receptor-specific target genes only harboring nearby binding sites that are shared (Fig. 2D, (Arora et al. 2013)). Here, we present evidence that shared binding sites also play a role in directing functional diversification among paralogous TFs. This raises the question: How can genomic loci that are occupied by both AR and GR direct receptor-specific transcription responses? Based on our studies, the explanation is that the downstream consequences of binding differ between AR and GR in several ways. For instance, GR binding induces robust changes in H3K27ac levels whereas for AR the increase is much more subtle (Fig. 3B). In addition, shared binding sites show a receptor-specific potential to activate transcription in reporter assays and based on our genome editing studies for the *AQP3* and *GILZ* genes (Fig. 5C, Fig. 6). The different downstream consequences of binding could in turn be a result of receptor-specific interactions with cofactors, *e.g.* the mediator complex, as observed in our RIME assays. Notably, cofactors display specificity for distinct types of core promoters indicating that ‘compatibilities’ between cofactors and promoters can influence if a recruited cofactor influences gene expression or not (Haberle et al. 2019). Moreover, depending on the target gene examined, different cofactors are required for GR-dependent regulation (Stallcup and Poulard 2020; Chen et al. 2006; Sacta et al. 2018). This raises the possibility that for each gene different cofactors address the rate-limiting step of the Pol 2 transcription cycle for gene activation. In this scenario, receptor-specific gene activation from shared binding sites would occur when different cofactors are recruited by AR and GR.

In summary, our study suggests that both divergence in genomic occupancy and diversity in the events that occur downstream of binding contribute to functional diversification among TF paralogs (Fig. 9).

**Fig. 9.**
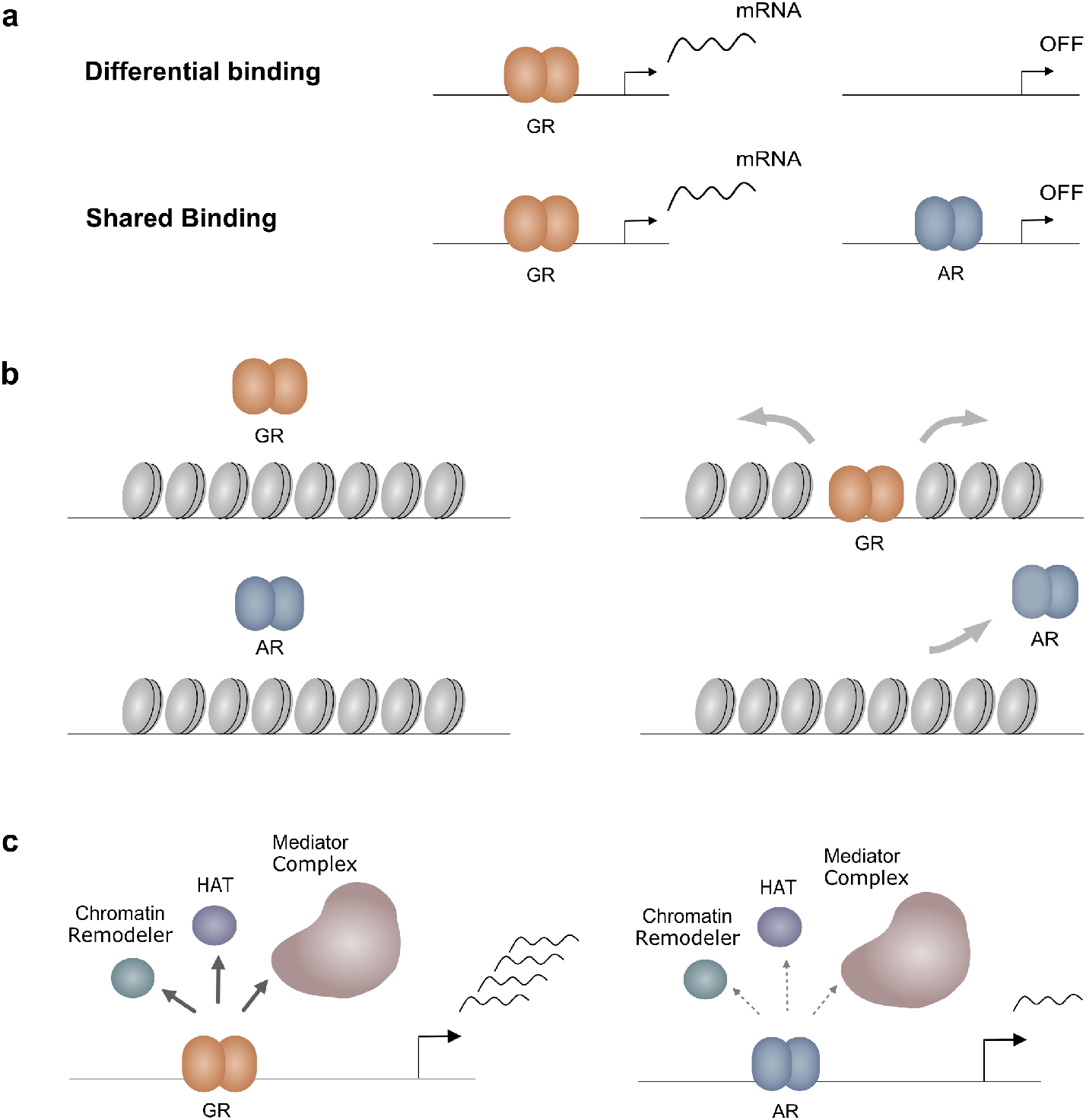
Models explaining receptor-specific regulation. (a) Specificity of transcriptional regulation can be driven by both receptor-specific binding and by selective activation from binding sites that are occupied by both AR and GR. (b) Receptor-specific binding due to a GR-specific ability to bind relatively inaccessible chromatin. (c) Receptor-specific interactions with cofactors can drive receptor-specific regulation from shared binding sites.

## Acknowledgements

We thank Edda Einfeldt for excellent technical support and Stefan Haas for help setting up the pipeline for RNA-seq analysis. This work was supported by the Else Kröner-Fresenius-Stiftung (grant 2014_A152 to S.H.M. and Me.B.).

## Author contributions

Ma.B. and S.H.M. performed and conceived the experiments and analysed the data. Me.B and G.K. analysed the data. A.F. and S.S. prepared the FAIRE-STARR library and established the assay. S.P. and I.M.P. performed and analysed the RIME experiments. C.H. and F.C. provided reagents. H.C, M.V. and W.Z supervised the study. S.H.M. wrote the manuscript with input from all authors.

---

**Fig. S1.**
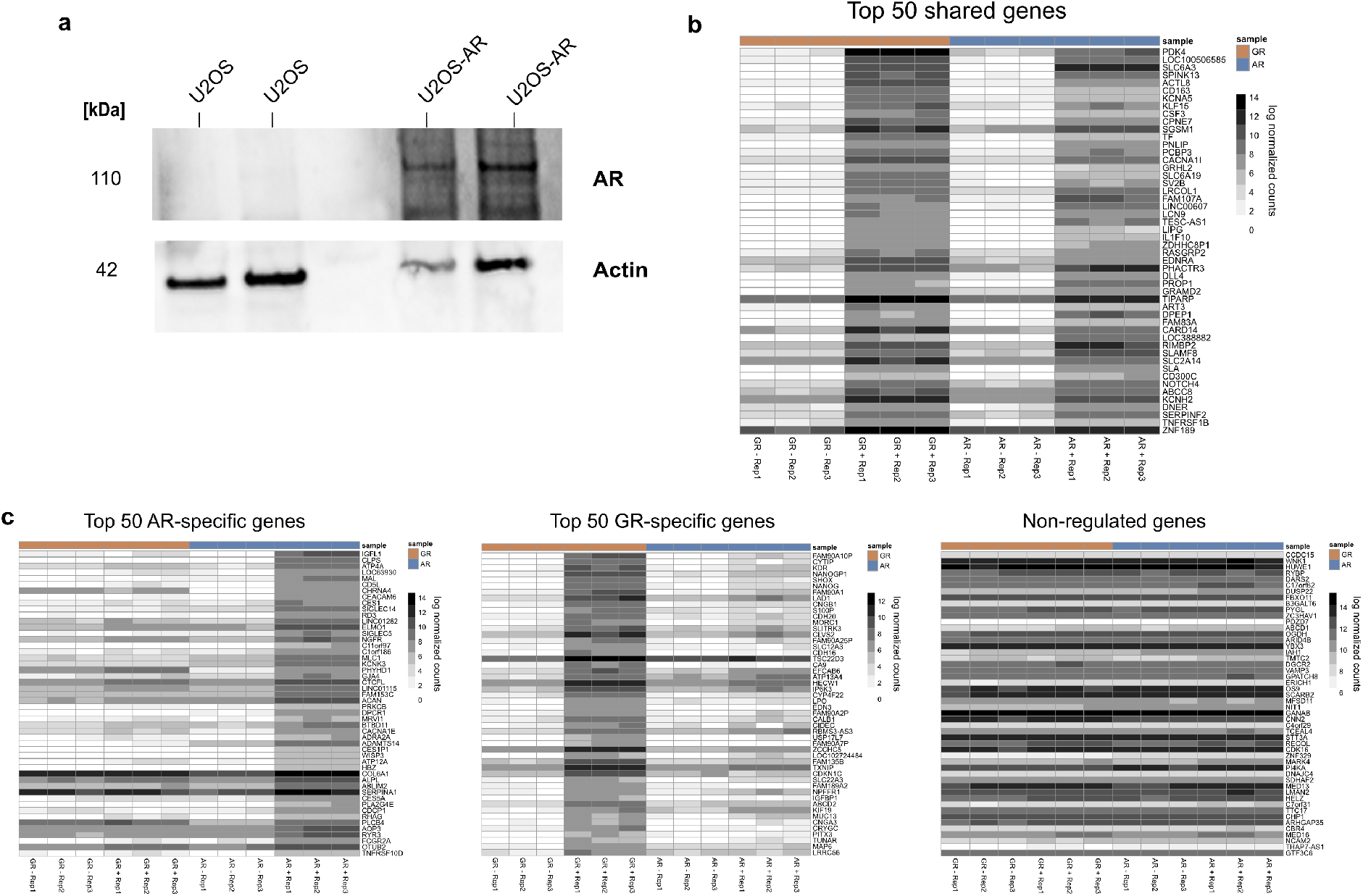
Analysis of the U2OS-AR cell line and heatmaps of different gene classes. (a) Total cell extracts of parental U2OS cells and a single-cell-derived U2OS-AR line were probed by Western blotting for the expression of AR and actin as loading control. (b) Heatmap displaying log normalized gene expression for the top 50 shared target genes (adjusted p-value < 0.05 and log2(fold change) > 1.5). Each column represents an individual replicate sample of either AR or GR (GR -, GR +, AR -, AR+), with rows representing individual genes. U2OS-AR cells were treated for 24h with 5 nM R1881; U2OS-GR cells were treated for 4h with 1 μM Dex. (c) Same as (b) except that top 50 AR-specific genes are shown. (d) Same as (b) except that top 50 GR-specific genes are shown. (e) Same as (b) except that 50 randomly selected non-regulated genes (adjusted p-value < 0.5 and 0.5 > log2(fold change) > 0) are shown.

**Fig. S2.**
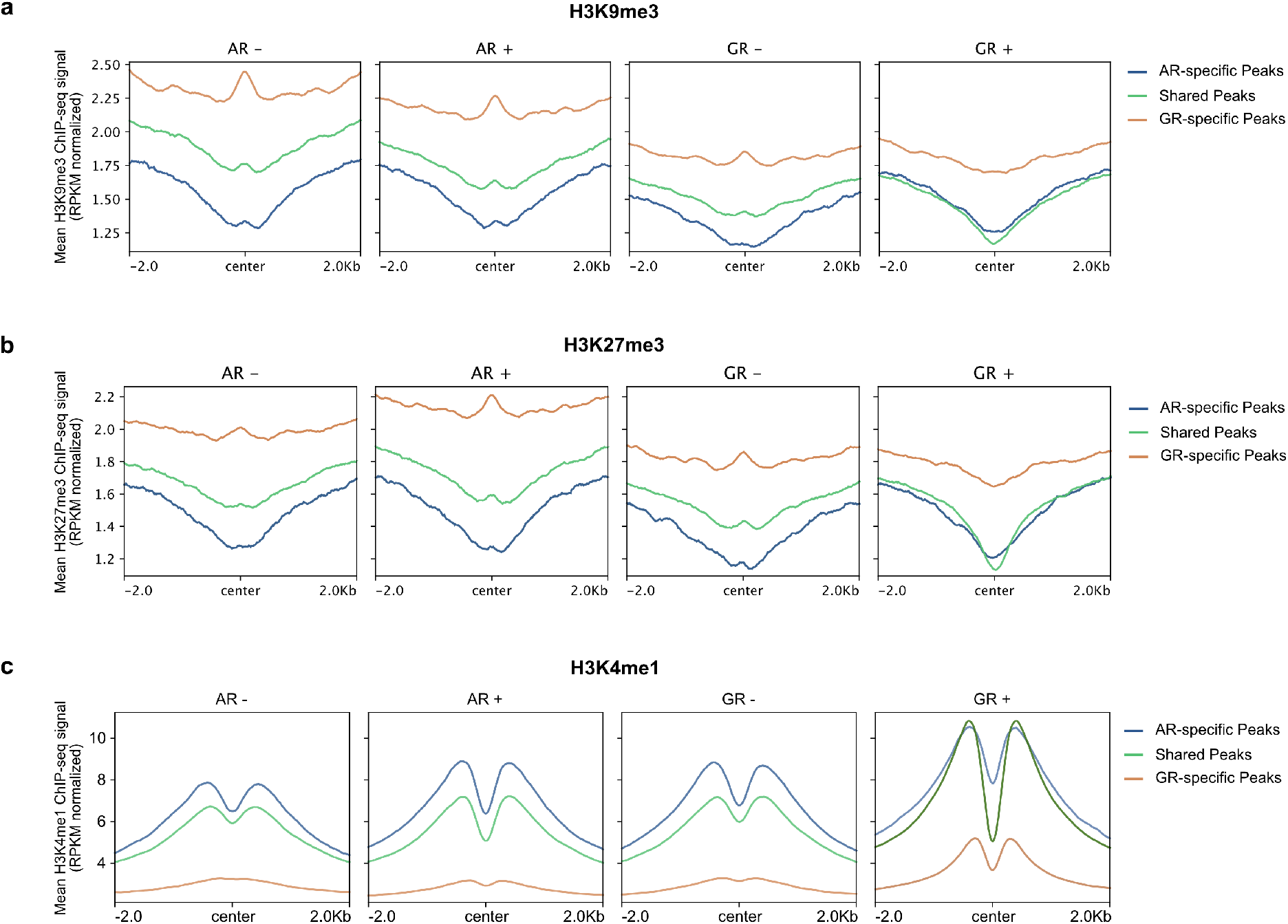
Mean signal plots for chromatin marks at shared and receptor-specific peaks. Mean signal plot of (a) H3K9me3, (b) H3K27me3 and (c) H3K4me1 ChIP-seq read coverage (RPKM normalized) at shared and receptor-specific binding sites (+/− 2 kb around peak center). U2OS-AR cells were treated with R1881 or vehicle (5 nM, 4h) and U2OS-GR cells with Dex or vehicle (1 μM, 1.5h).

**Fig. S3.**
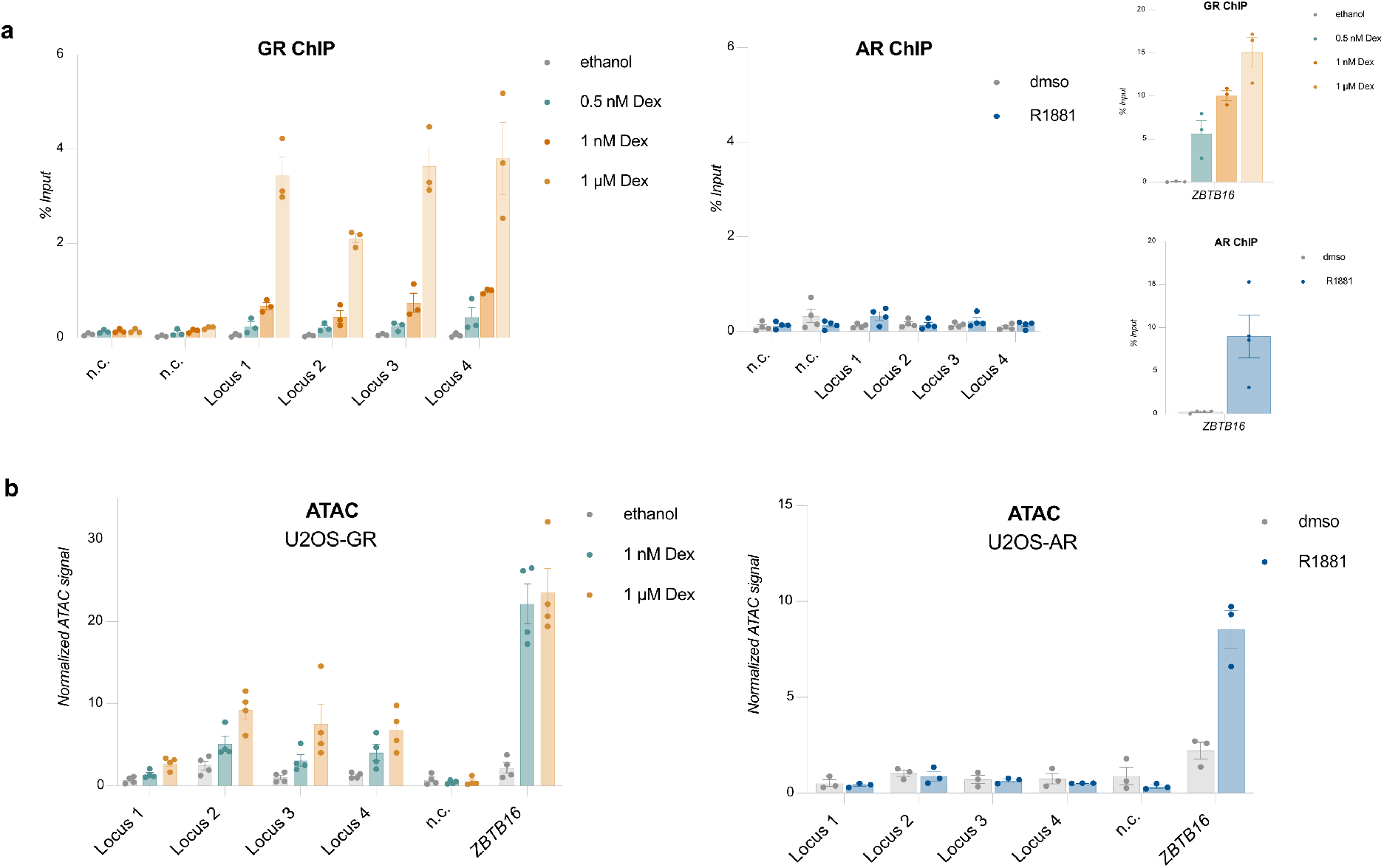
GR binding and ATAC signal at low hormone concentrations. (a) ChIP-qPCR of GR-specific peaks as indicated in U2OS-GR and U2OS-AR cells. U2OS-GR cells were treated with 0.5 nM, 1 nM, 1 μM Dex or ethanol as a vehicle control for 1.5h. U2OS-AR cells were treated with 5 nM R1881 or dmso for 4h. Average percentage of input precipitated ± SEM is shown (n ≥ 3). n.c.: negative control region Right: binding of either GR or AR at a shared binding site near the *ZBTB16* gene. (b) Same as for (a) except that ATAC-qPCR signal normalized to genomic DNA is shown.

**Fig. S4.**
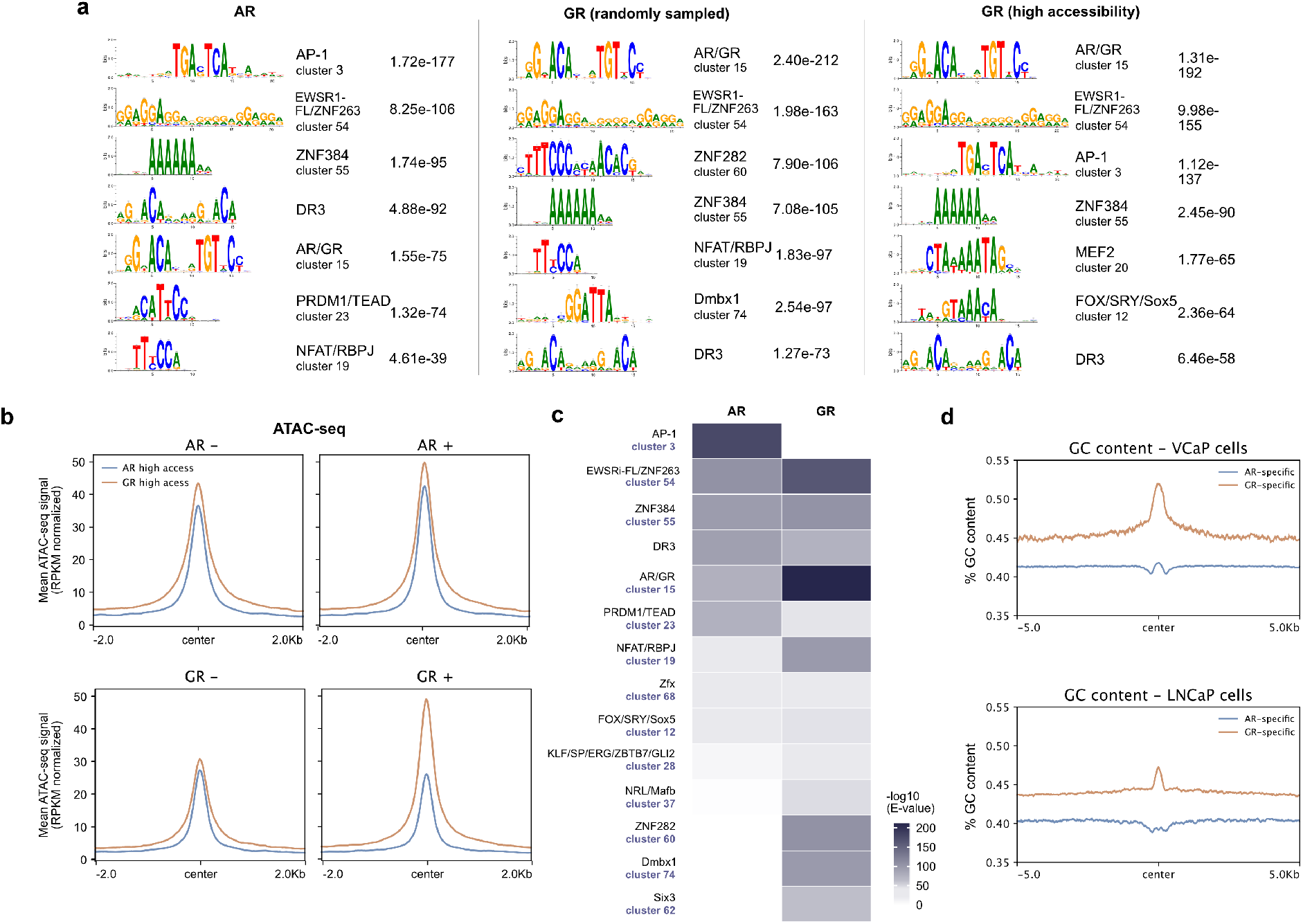
Sequence motif analysis. (a) Top 7 enriched motif clusters and corresponding E values at all AR-specific peaks, an equal number of randomly sampled GR-specific peaks or GR-specific peaks in regions with high chromatin accessibility (+/− 250 pb around the peak center). Shuffled input sequences were used as background for the motif enrichment analysis. (b) Mean signal plots of ATAC-seq read coverage (RPKM normalized) at high accessibility AR- and GR-specific sites as used in Fig. 4C (+/− 2 kb around peak center). U2OS-AR cells were treated with vehicle or R1881 (5 nM, 4h) and U2OS-GR cells with vehicle or Dex (1 μM, 1.5h).(c) Heatmap visualization of enriched motif clusters (from JASPAR 2018 CORE Vertebrates Clustering motifs) at all AR-specific or at an equal number of randomly sampled GR-specific peaks (+/− 250 pb around the peak center). Shuffled input sequences were used as background for the motif enrichment analysis. Motifs were included if the E value was <10-30 for either AR or GR. (d) GC content at GR- and AR-specifically occupied regions in VCaP and LNCaP cells (+/− 5 kb around the peak center).

**Fig. S5.**
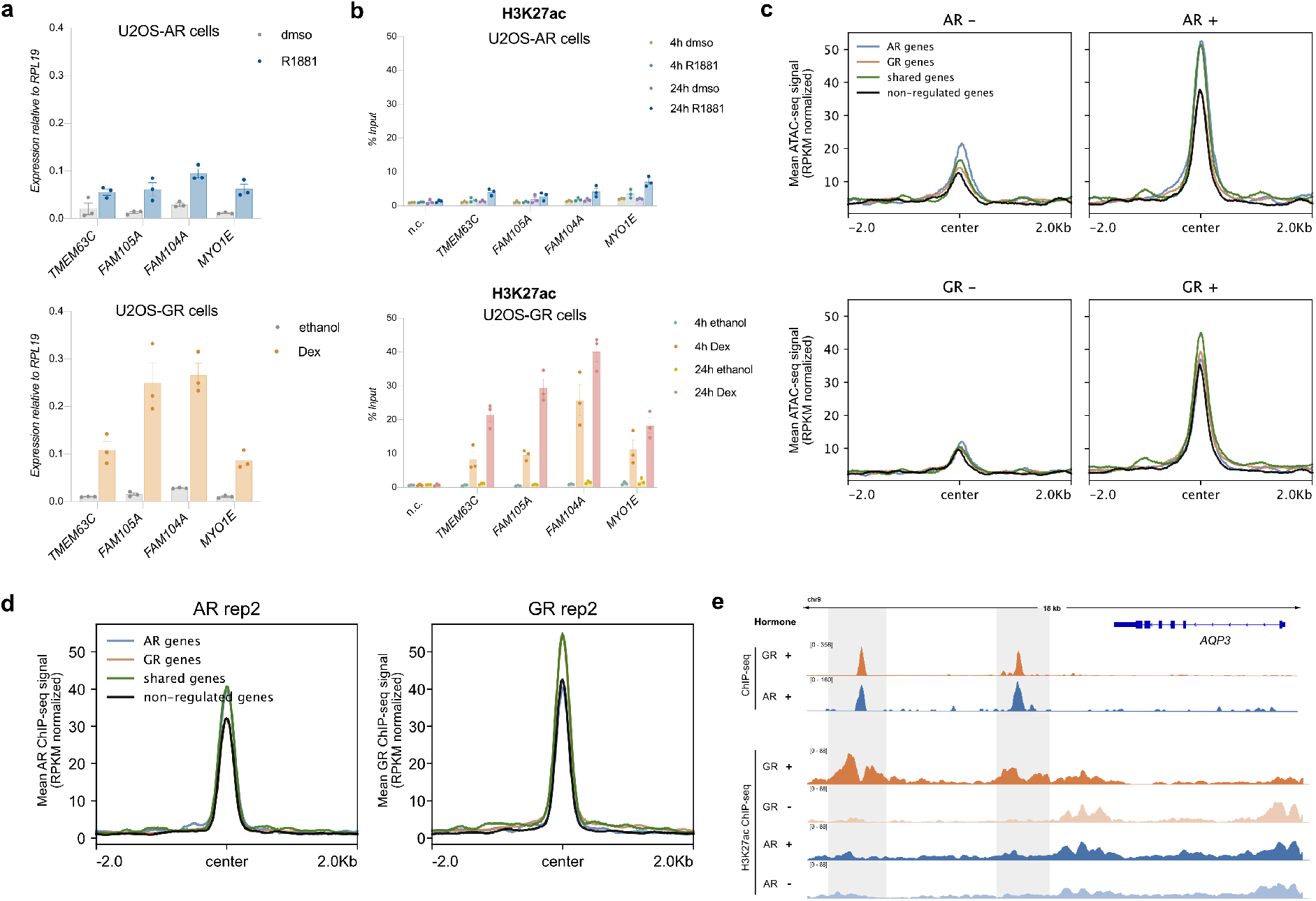
Shared regulation, receptor and H3K27ac ChIP-seq and ATAC-seq at shared binding sites. (a) Relative mRNA levels of *TMEM63C*, *FAM105A*, *FAM104A* and *MYO1E* were quantified by qPCR for U2OS cells stably expressing either (top) AR or bottom (GR). U2OS-AR cells were treated for 24h with dmso as vehicle control or 5 nM R1881. U2OS-GR cells with ethanol as vehicle control or 1 μM Dex for 24h. Average gene expression ±SEM is shown (n = 3). (b) H3K27ac levels were analyzed by ChIP for regions occupied by both AR and GR near the shared target genes as indicated. U2OS-AR cells were treated for either 4h or 24h with dmso as vehicle control or 5 nM R1881. U2OS-GR cells were treated for either 4h or 24h with ethanol as vehicle control or 1 μM Dex. Average percentage of input precipitated ± SEM is shown (n = 3). n.c.: negative control region. (c) Mean signal plot of ATAC-seq read coverage (RPKM normalized) at shared sites (+/− 2 kb around the peak center) near the different gene categories as used in Fig. 5C. U2OS-AR cells were treated with R1881 or vehicle (5 nM, 4h) and U2OS-GR cells with Dex or vehicle (1 μM, 1.5h). (d) Same as (c) except that AR and GR ChIP-seq read coverage (RPKM normalized) is shown. (e) Genome browser screenshot for the *AQP3* locus showing receptor and H3K27ac ChIP-seq signals for cells treated as indicated. Regions bound by both receptors are highlighted in grey.

**Fig. S6.**
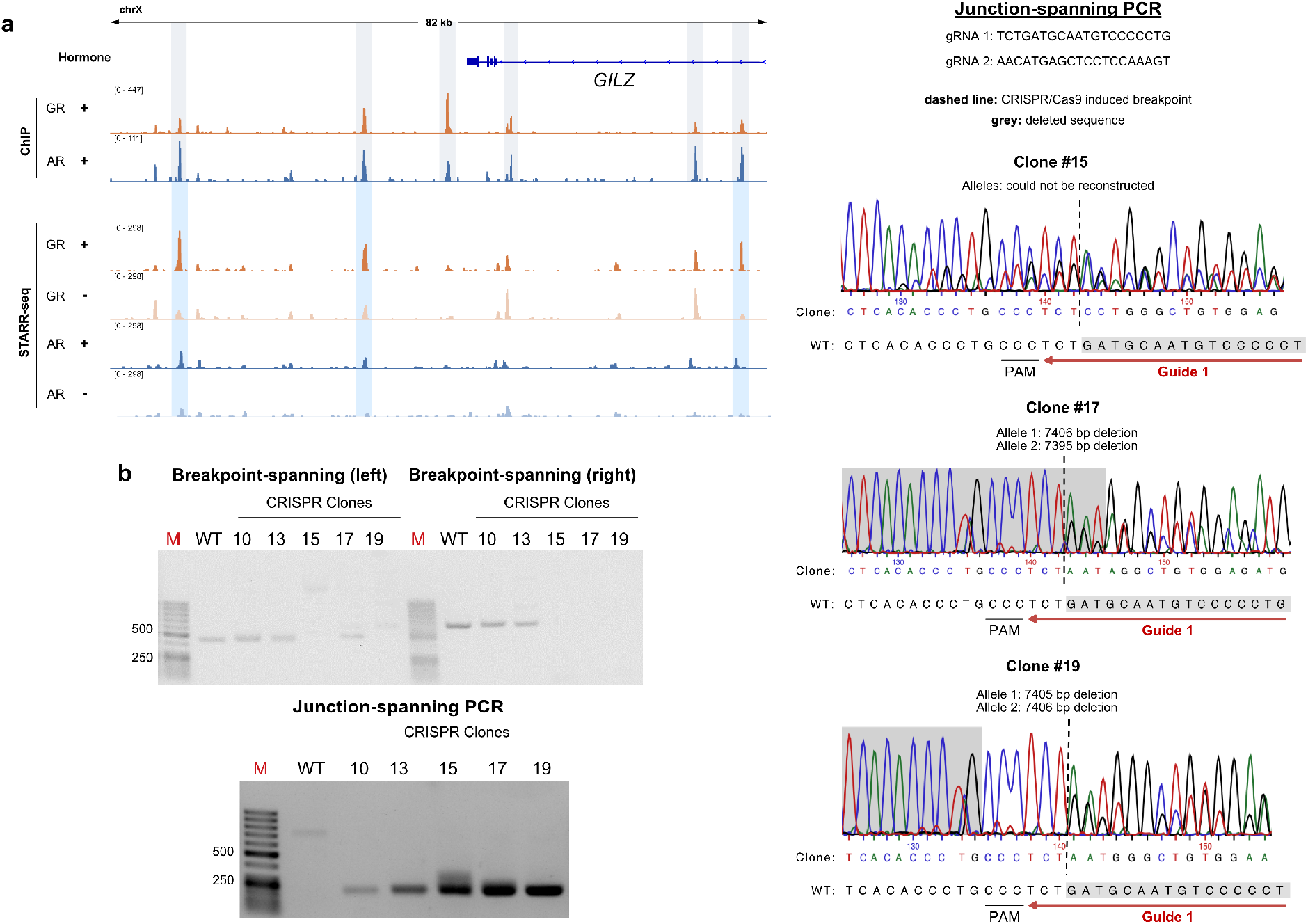
Receptor binding and FAIRE-STARR signal at the *GILZ* locus/genotyping of clonal lines. (a) Genome browser screenshot of the *GILZ* locus showing ChIP-seq and STARR-FAIRE-seq tracks for GR and AR. The STARR-seq tracks show enhancer activity in U2OS-GR cells treated with ethanol or 1 μM Dex and U2OS-AR cells treated with dmso or 5 nM R1881 overnight. The receptor-bound peaks are highlighted in grey and hormone-inducible STARR-seq enhancers are highlighted in blue. STARR-seq tracks depict the merged signal from three biological replicates. (b) Left: To genotype single-cell-derived U2OS-AR clonal lines, the CRISPR-targeted *AQP3* locus was PCR amplified from genomic DNA and analyzed on agarose gels. For the junction-PCR, primers are placed outside the breakpoints to detect CRISPR clones carrying a genomic deletion at the *AQP3* locus. Primers flanking the breakpoints were used to detect the presence of WT alleles. The expected amplicon size of the left WT breakpoint is 476 bp and 686 bp for the right WT breakpoint. For clone #17, a signal is observed for the left WT breakpoint of clone #17 indicating a partial deletion of the second allele. M represents the DNA size marker GeneRuler 50bp. Right: Sanger sequencing results of the junction-PCR amplicon for successfully edited clonal lines #15, #17 and #19 and the inferred deletion for each allele.

**Fig. S7.**
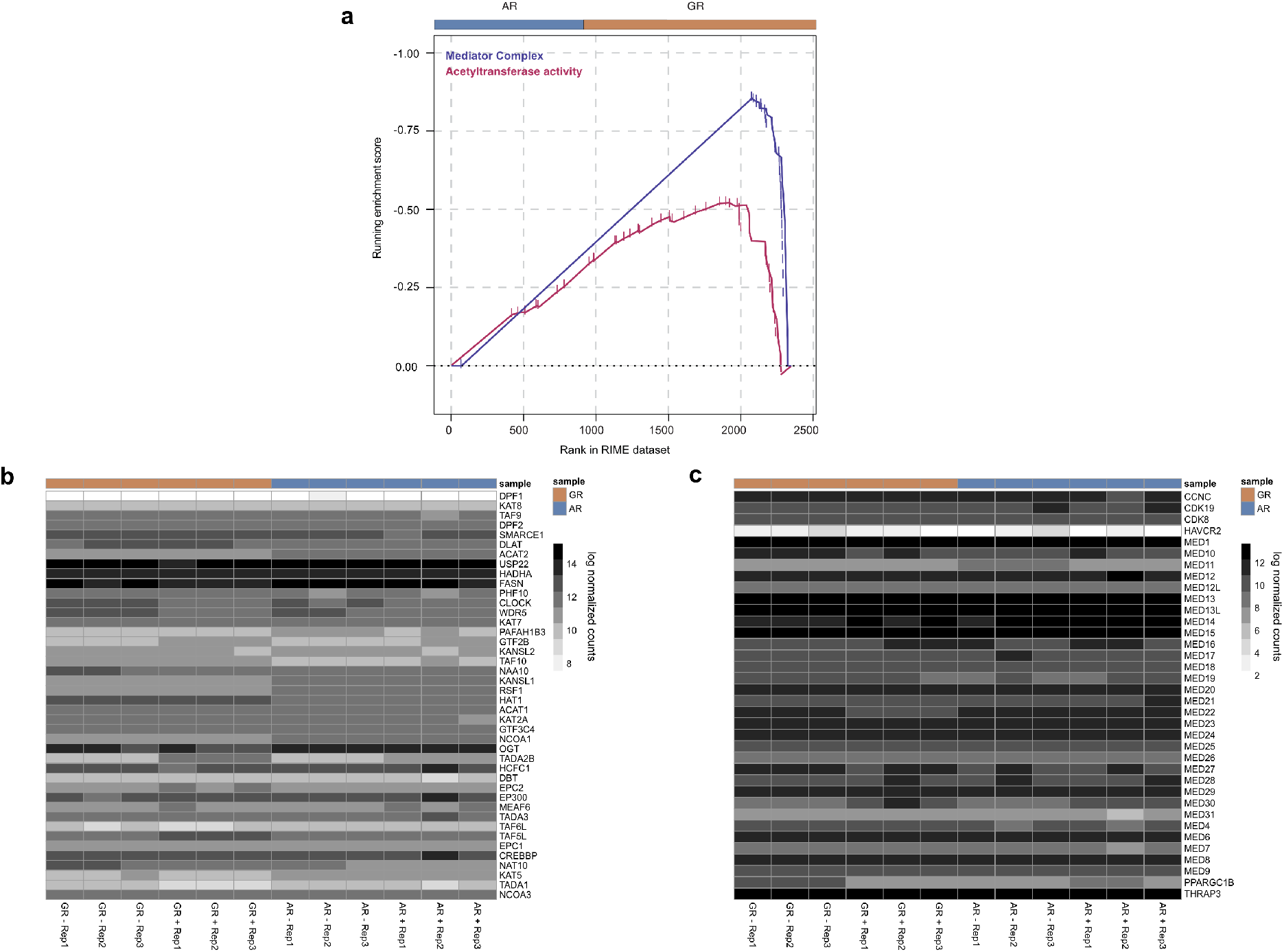
Gene Set Enrichment Analysis of the RIME data. (a) Gene Set Enrichment Analysis (GSEA) results comparing the RIME data for AR vs GR. Two pathways with GR-specific enrichment are shown. (b) Heatmap displaying log normalized gene expression for proteins from the acetyltransferase activity category that selectively interacted with GR. U2OS-AR cells were treated for 24h with 5 nM R1881; U2OS-GR cells for 4h with 1 μM Dex. (c) Same as for (b) except for genes of the mediator complex category.

**Figure S8.**
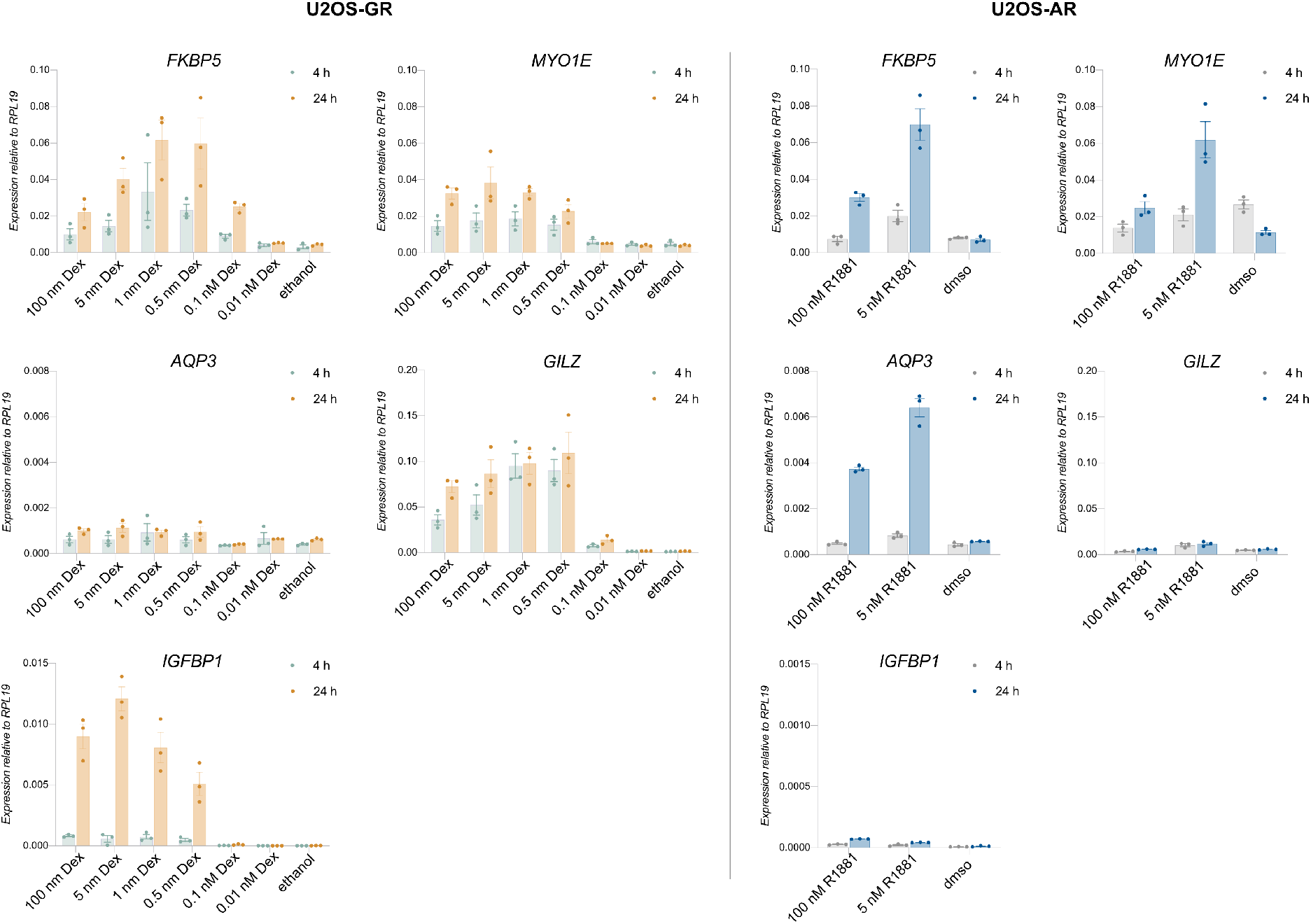
Gene regulation at different hormone concentrations and time points. (a) Relative mRNA levels of genes as indicated was quantified by qPCR for U2OS cells stably expressing either (left) GR or (right) AR. U2OS-AR cells were treated for 4h or 24h with dmso as vehicle control or with R1881 concentration as indicated. U2OS-GR cells were treated with ethanol as vehicle control or with Dex concentration as indicated for 4h or 24h. Average gene expression ±SEM is shown (n = 3).

**Figure S9.**
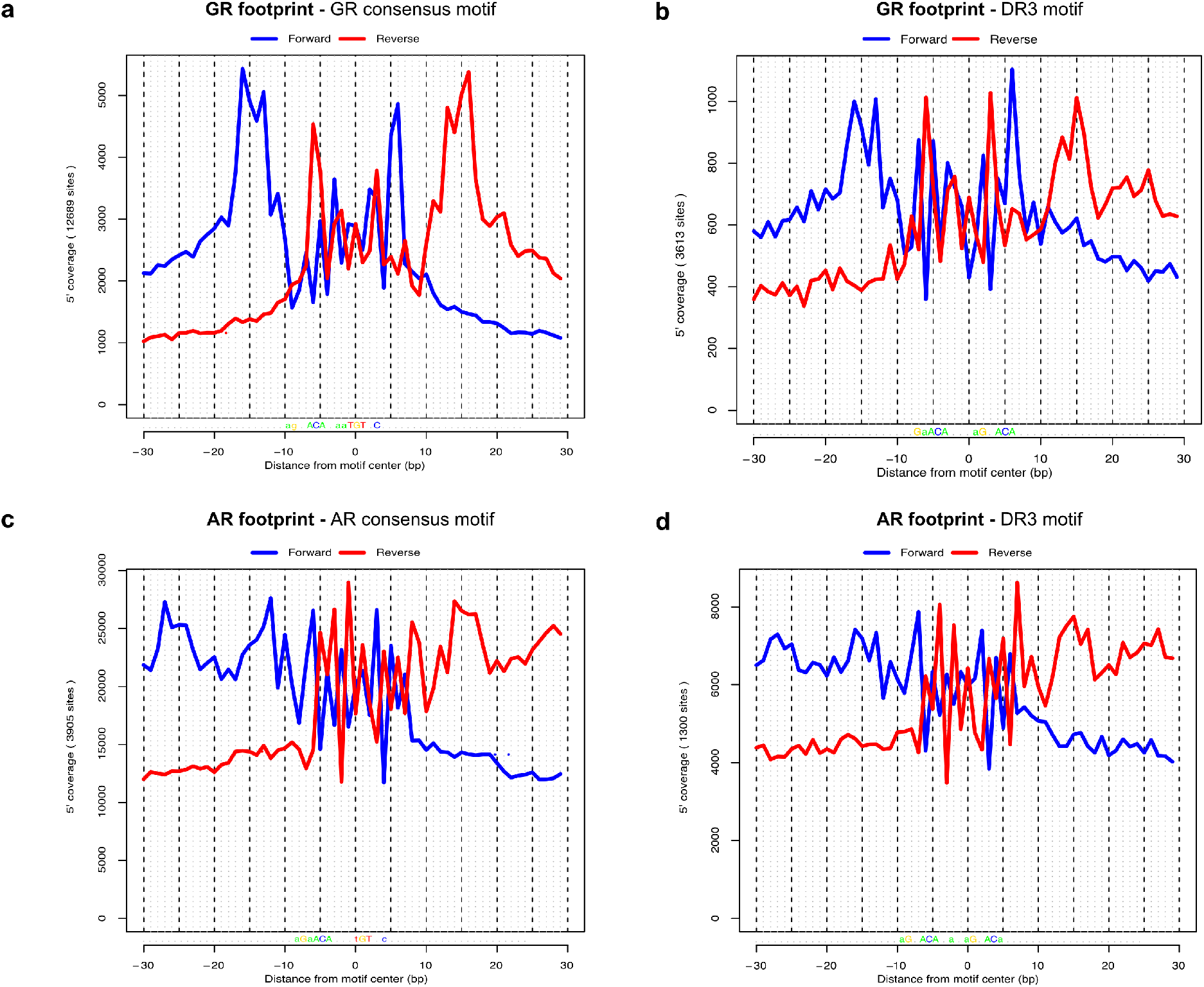
ChIP-exo profiles for AR and GR at canonical and at direct repeat-like (DR3) sequences. (a) GR ChIP-exo footprint profile for the GR consensus motif in U2OS-GR cells. Blue represents ChIP-exo signal for the positive and red the negative strand. (b) GR ChIP-exo footprint profile for the DR3 motif in U2OS-GR cells. (c) AR ChIP-exo footprint profile for the AR consensus motif in LNCaP cells. (d) AR ChIP-exo footprint profile for the DR3 motif in LNCaP cells.

